# SUMOylation of SAMHD1 at Lysine 595 is required for HIV-1 restriction in non-cycling cells

**DOI:** 10.1101/2020.06.04.133439

**Authors:** Charlotte Martinat, Arthur Cormier, Joёlle Tobaly-Tapiero, Noé Palmic, Nicoletta Casartelli, Si’Ana A. Coggins, Julian Buchrieser, Mirjana Persaud, Felipe Diaz-Griffero, Lucile Espert, Guillaume Bossis, Pascale Lesage, Olivier Schwartz, Baek Kim, Florence Margottin-Goguet, Ali Saïb, Alessia Zamborlini

## Abstract

SAMHD1 is a cellular triphosphohydrolase (dNTPase) proposed to inhibit HIV-1 reverse transcription in non-cycling immune cells by limiting the supply of the dNTP substrates. Yet, phosphorylation of T592 downregulates SAMHD1 antiviral activity, but not its dNTPase function, implying that additional mechanisms contribute to viral restriction. Here, we show that SAMHD1 is SUMOylated on residue K595, a modification that relies on the presence of a proximal SUMO-interacting motif (SIM). Loss of K595 SUMOylation suppresses the restriction activity of SAMHD1, even in the context of the constitutively active phospho-ablative T592A mutant but has no impact on dNTP depletion. Conversely, the artificial fusion of SUMO to a non-SUMOylatable inactive SAMHD1 variant restores its antiviral function. These observations clearly establish that the absence of T592 phosphorylation cannot fully account for the restriction activity of SAMHD1. We find that concomitant SUMOylation of K595 is required to stimulate a dNTPase-independent antiviral activity.

## Introduction

Sterile alpha-motif (SAM) and histidine-aspartate (HD) domain-containing protein 1 (SAMHD1) is a cellular triphosphohydrolase (dNTPase) that inhibits the replication of the human immunodeficiency virus type 1 (HIV-1) in non-cycling immune cells such as macrophages, monocytes, dendritic cells and resting T4 lymphocytes^1–5^. This antiviral function is largely attributed to the ability of SAMHD1 to hydrolyze dNTPs into the desoxynucleoside and triphosphate components^6–8^ thereby reducing the cellular dNTP supply below a threshold required for efficient reverse transcription of the viral RNA genome^9,10^. In contrast to HIV-1, the related HIV-2 virus counteracts this restriction by expressing the Vpx accessory protein which promotes the degradation of SAMHD1 through the ubiquitin-proteasome system^3–5,11,12^. SAMHD1 depletion is accompanied by both dNTP pools expansion and increased cell permissiveness to HIV-1 infection^8,13^ indicating that the dNTPase and restriction functions are linked.

SAMHD1 is an ubiquitous protein^2,14,15^. Yet, its anti-HIV-1 activity is witnessed only in non-cycling cells, pointing to the involvement of post-translational regulatory mechanisms. It is now well established that residue T592 is phosphorylated by the cyclin/CDK complexes during the G1/S transition^16–18^ and dephosphorylated by members of the phosphoprotein phosphatase (PPP) family upon mitotic exit^19,20^. This modification likely enables SAMHD1 to promote the progression of replication forks in dividing cells^21^. Phosphorylation at T592 is weak to undetectable in non-cycling cells refractory to HIV-1 ^19,20,22^, suggesting that only dephosphorylated SAMHD1 might be restriction-competent. Consistent with this model, mutation of T592 into D or E to mimic phosphorylation renders SAMHD1 antivirally inactive^17,18,23,24^. However, the phosphomimetic variants retain WT dNTPase function^18,24,25^.

In the same line, SAMHD1 prevents dNTP pools expansion throughout the cell cycle, regardless of its phosphorylation status^20^. Altogether these data question whether the establishment of a SAMHD1-mediated antiviral state might only rely on dNTP depletion and/or regulation by T592 phosphorylation. Reports that SAMHD1 degrades the incoming viral RNA genome through a ribonuclease activity remain controversial^23,26–30^, calling for additional investigations to clarify the molecular mechanisms underlying its viral restriction function.

Interestingly, SAMHD1 was a hit in recent large-scale proteomic studies investigating the cellular substrates of SUMOylation^31,32^, a dynamic post-translational modification (PTM) and an important regulator of many fundamental cellular processes including immune responses^33^. SUMOylation consists in the conjugation of a single Small Ubiquitin-like Modifier (SUMO) moiety or a polymeric SUMO chain to a protein substrate through the sequential action of a dedicated set of E1-activating, E2-conjugating and E3-ligating enzymes. SUMOylation is reversed by SUMO-specific proteases (e.g. SENP)^34^. Human cells express three ubiquitous SUMO paralogs. SUMO1 shares ~50% of sequence homology with SUMO2 and SUMO3, which are ~90% similar and thus referred to as SUMO2/3^35^. SUMOylation often targets the Lysine (K) residue lying within the consensus motif φKxα (φ: hydrophobic amino acid, x: any amino acid and α: an acidic residue) that represents the binding site for the unique SUMO E2 conjugating enzyme Ubc9^36,37^. A proximal SUMO-interacting motif (SIM), which typically consists of a short stretch of surface-exposed aliphatic residues^38^, might sometimes contribute to the recruitment and the optimal orientation of the SUMO-charged Ubc9, allowing an efficient transfer of SUMO to the substrate. A SIM might also constitute a binding interface for SUMO-conjugated partners that mediate the downstream consequences of SUMOylation.

In this study we show that SAMHD1 undergoes SIM-mediated SUMOylation of the evolutionarily conserved K595 residue, which is part of the CDK-targeted motif driving T592 phosphorylation (^592^TPQK^595^). Preventing K595 SUMOylation by mutation of either key residues of the SUMO-consensus motif or the SIM (^488^LLDV^501^) invariably suppressed SAMHD1-mediated viral restriction, but not its dNTPase activity. This was true even when T592 was dephosphorylated and therefore SAMHD1 expected to be antivirally active. These observations suggest that the status of T592 phosphorylation cannot fully account for the regulation of the restriction activity of SAMHD1. Finding that the artificial fusion of SUMO to an inactive C-terminal truncation mutant (lacking both T592 phosphorylation and K595 SUMOylation) restored the inhibition of viral infection, further supports the requirement of SUMO conjugation to K595 for the establishment of a SAMHD1-dependent antiviral state in non-cycling immune cells.

## Results

### SAMHD1 is a SUMO target

To investigate if SAMHD1 is a SUMO substrate, we used a 293T cell-based assay where we expressed HA-SAMHD1 together with each 6xHis-tagged SUMO paralog and the SUMO E2 conjugating enzyme Ubc9. Cells were treated or not with the proteasome inhibitor MG132, to favor the accumulation of SUMO-conjugated proteins^39^. Next, samples were lysed in denaturing conditions to inhibit the highly active SUMO proteases and preserve SUMOylation. Following enrichment of SUMO-conjugated proteins by histidine affinity, the fraction of SUMOylated SAMHD1 was detected with an anti-HA antibody. A ~100 kDa band, consistent with the expected size of SAMHD1 conjugated to a SUMO moiety, was visualized in SUMO2- and SUMO3-expressing cells, at baseline (**Fig. 1A**, lanes 3 and 4). Proteasome inhibition caused an accumulation of high molecular-weight SAMHD1 species, which are likely SUMO chain conjugates, pointing to a potential role of SUMOylation in the control of the protein turnover (**Fig. 1A**, lanes 7 and 8). Modified SAMHD1 forms were undetectable in cells expressing SUMO1 or transfected with the empty plasmid, although the expression levels of SAMHD1 and SUMO isoforms were similar in all samples (**Fig. 1A**, lanes 1, 2, 5 and 6).

**Figure 1.**
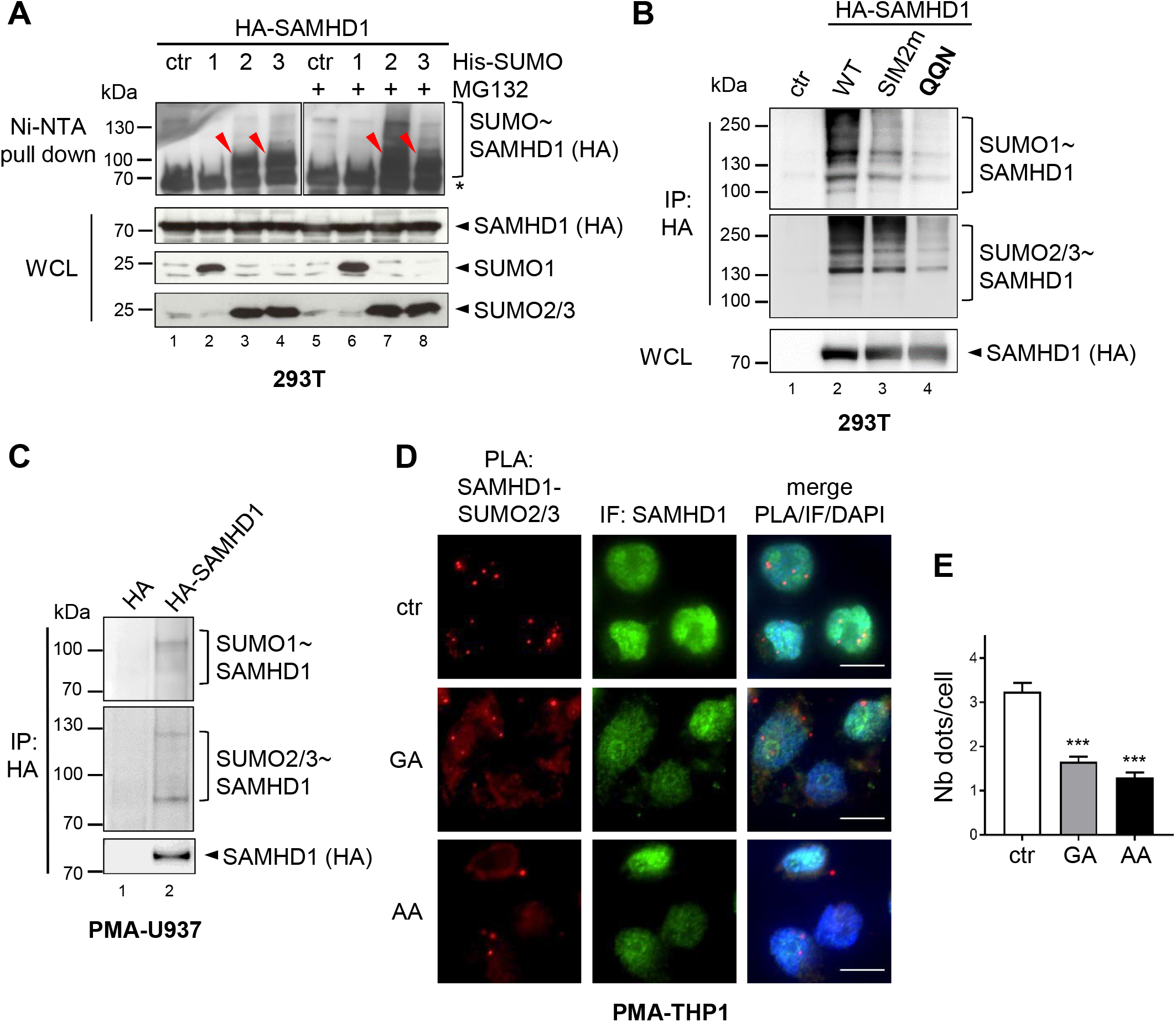
SAMHD1 is SUMOylated both in dividing and non-dividing cells. **A.** HEK 293T cells were transfected with plasmids encoding HA-SAMHD1 and N-terminal His-tagged SUMO1 (1), SUMO2 (2), SUMO3 (3) or the control empty plasmid (ctr). After 48 hours, cells were lyzed in denaturing conditions (guanidium HCl 6M, imidazole 10 mM) to inhibit SUMO proteases, and SUMO-conjugates were enriched on Ni-NTA beads. Proteins contained in the eluates or the whole cell lysates (WCL) were separated on a 4-15% SDS-PAGE gel and detected by immunoblotting using anti-HA or anti-SUMO paralog specific antibodies. A representative result is shown (n=3). Arrowheads point to SAMHD1~SUMO conjugates. *, nonspecific binding of unmodified SAMHD1 on Ni-NTA beads. **B.** HEK 293T cells transiently expressing HA-SAMHD1 WT or SUMOylation-deficient SIM2m and **QQN** variants were lysed in buffer containing 1% SDS. After dilution, SAMHD1 and its post-translational derivatives were immunoprecipitated on HA-matrix beads and analyzed as in A. A representative result is shown (n=2). **C.** U937 cells stably expressing HA-SAMHD1 or the empty vector control (HA) were induced to differentiate into macrophage-like cells by treatment with PMA (100 ng/mL, 24 h). SAMHD1 SUMOylation was assessed as in B. **D.** Differentiated THP-1 cells were incubated with gingkolic acid (GA, 50μM), anacardic acid (AA, 50μM) or DMSO (ctr) for 2 hours. After fixation, cells were probed with anti-SAMHD1 and anti-SUMO2/3 antibodies before being processed for Proximity Ligation Assay (PLA panels, red dots). SAMHD1 localization was analyzed using an anti-isotype secondary antibody coupled to the Alexa_488_ dye (IF panels, green). Nuclei were stained with DAPI. Scale bar = 10 μm. Representative images are shown (n=2) **E.** The PLA signal was quantified manually using the same thresholding parameters across parallel samples as defined with the Icy software. Bars represent the average number of dots per cell ±SD (n=150). Significance was determined by one-way ANOVA statistical test. ***: p < 0.001.

Having confirmed that SAMHD1 is modified by ectopically expressed SUMO paralogs, we tested its conjugation by endogenous SUMOs. To this aim, HA-SAMHD1 was over-expressed by either transfection of 293T cells or lentiviral transduction of the human monocytic U937 cell line, which lacks measurable levels of endogenous SAMHD1 and acquires a macrophage-like phenotype when exposed to phorbol 12-myristate 13-acetate (PMA). After lysis in stringent conditions, SAMHD1 and its post-translational derivatives were enriched on HA-matrix beads. Consistent with the previous experiment, a ladder of SUMO-conjugates was visualized with anti-SUMO2/3 antibodies in both actively dividing (**Fig. 1B**, 293T) and non-dividing cells (**Fig. 1C**, PMA-U937). Under these experimental conditions, modification of SAMHD1 by SUMO1 was also detected (**Fig. 1B** and **1C**, upper panels, lane 2).

To extend these findings, we used the proximity-ligation assay (PLA)^40,41^ to analyze the interaction between endogenous SAMHD1 and SUMO in human monocytic THP1 cells differentiated into macrophage-like cells by PMA treatment. As both SAMHD1 and the SUMO machinery are enriched in the nucleus, it was not surprising to detect fluorescent dots (~3.24±0.2, per cell) indicative of the SAMHD1-SUMO2/3 association mainly in this compartment (**Fig. 1D**). Importantly, a 2-hour incubation with either ginkgolic acid (GA) or its structurally related analog anacardic acid (AA), which block SUMO conjugation by inhibiting the E1 SUMO-activating enzyme^42^, lowered the frequency of the PLA signal by ~2- to ~2.8-fold, respectively, thereby confirming the relevance of the PLA approach to study the SAMHD1-SUMO2/3 interaction (**Fig. 1D**). While the proximity labeling was markedly diminished upon exposure to SUMOylation inhibitors, SAMHD1 localization (**Fig. 1D**) and general expression as well as the global amount of SUMO2/3-conjugates were unaffected (**Fig. S1**). An interaction between SAMHD1 and SUMO1 was also visualized in the nucleus of differentiated THP1, but not SAMHD1-negative U937 cells, further validating the specificity of the SAMHD1-SUMO PLA signal (**Fig. S2**). Altogether, these data show that SAMHD1 is conjugated by SUMO1 and SUMO2/3 in the nucleus of both cycling and differentiated cells.

### Lysine residues at position 469, 595 and 622 are the main SUMOylation sites of SAMHD1

Among several potential SUMO-attachment sites identified by high-resolution proteomic studies in human SAMHD1, residues K469, K595 and K622 were the most frequent hits (**Table S1**). Protein sequence alignment shows that the position corresponding to amino acid 595 of human SAMHD1, which is the last residue of the CDK-targeted ^592^TPQK^595^ motif (general consensus [S/T]Px[K/R]^43^), is invariably occupied by K, except for the murine isoform 2 (**Fig. S3**). Conversely, K469 and K622 are conserved among primate orthologs, with the former also found in prosimian, equine and koala isoforms (**Fig. S3**). To confirm that the identified sites are modified by SUMO, we performed the 293T-based SUMOylation assay using SAMHD1 mutants where the candidate K residues were changed into either arginine (R), to preserve a basic character, or alanine (A) (**Fig. 2A**). Alternatively, we mutated the acidic residue at position +2 of the SUMO-acceptor K residue that is essential for the recruitment of Ubc9, the unique E2 SUMO-conjugating enzyme^36,37^ (**Fig. 2A**). We focused our analyses on the SUMO2 paralog because i) the pool of SUMO2 and SUMO3 available for conjugation exceeds that of SUMO1^44^ and ii) SUMO2 and SUMO3 differ only by three amino acids and are undistinguishable with available antibodies^45^. Mutation of individual amino acids had a negligible effect on the electrophoretic mobility of SAMHD1 SUMO-conjugates both at baseline (**Fig. 2B**) and upon proteasome inhibition (**Fig. S4A**) indicating that multiple sites might be modified simultaneously. To test this idea, we monitored the SUMOylation pattern of SAMHD1 variants where candidate sites were inactivated in various combination. Hereafter, these mutants are named by a three-letter code corresponding to the residues found at position 469, 595 and 622 (where K was replaced by either R or A) or 471, 597 and 624 (where E or D were replaced by Q or N, respectively). The simultaneous K_595_A and K_622_R changes (yielding the K**AR** variant, where K at position 469 is intact) prevented the formation of the ~100 kDa band seen with WT SAMHD1 and likely representing a mono-SUMOylated form (**Fig. 2C**, lanes 2 and 3). This observation suggested that K595 and/or K622 are modified by a single SUMO moiety. Consistently, the mono-SUMOylated SAMHD1 form was detected only when the SUMO site centered on either K595 or K622 was intact (corresponding to mutants **R**K**R** and **Q**E**N** or **RA**K, respectively), although to a weaker extent relative to the WT protein (**Fig. 2B**, compare lanes 2 and 4, and lanes 7 and 8 to 6). We also analyzed the SUMOylation profile of SAMHD1 double mutants upon MG132 treatment. The K**AR** and **RA**K variants, where K469 or K622 is intact, respectively, displayed an altered polySUMOylation pattern as compared to WT SAMHD1 (**Fig. S4B**, compare lanes 3 and 4 to 2). Conversely, the simultaneous arginine substitution of K469 and K622 (yielding the **R**K**R** mutant) virtually abolished SAMHD1 polySUMOylation (**Fig. S4B**, compare lanes 7 and 8 to 6). The concomitant E471Q and D624N changes (yielding the **Q**E**N** mutant) had analogous consequences (**Fig. S4B**, compare lanes 6 and 8). These results indicate that K469 and K622, but not K595, are target sites for SUMO chains which accumulate upon MG132 treatment. Finally, we established that inactivation of the three SUMO-acceptor sites of SAMHD1 (**RAR** and **QQN** variants) strongly hampered the formation of slow migrating bands when SUMOs were expressed either ectopically (**Fig. 2C and S4B**, lanes 9 and 10) or at endogenous levels (**Fig. 1B**, compare lanes 2 and 4). Overall, these results confirm that residues K469, K595 and K622 are the major SUMOylation sites of SAMHD1.

**Figure 2.**
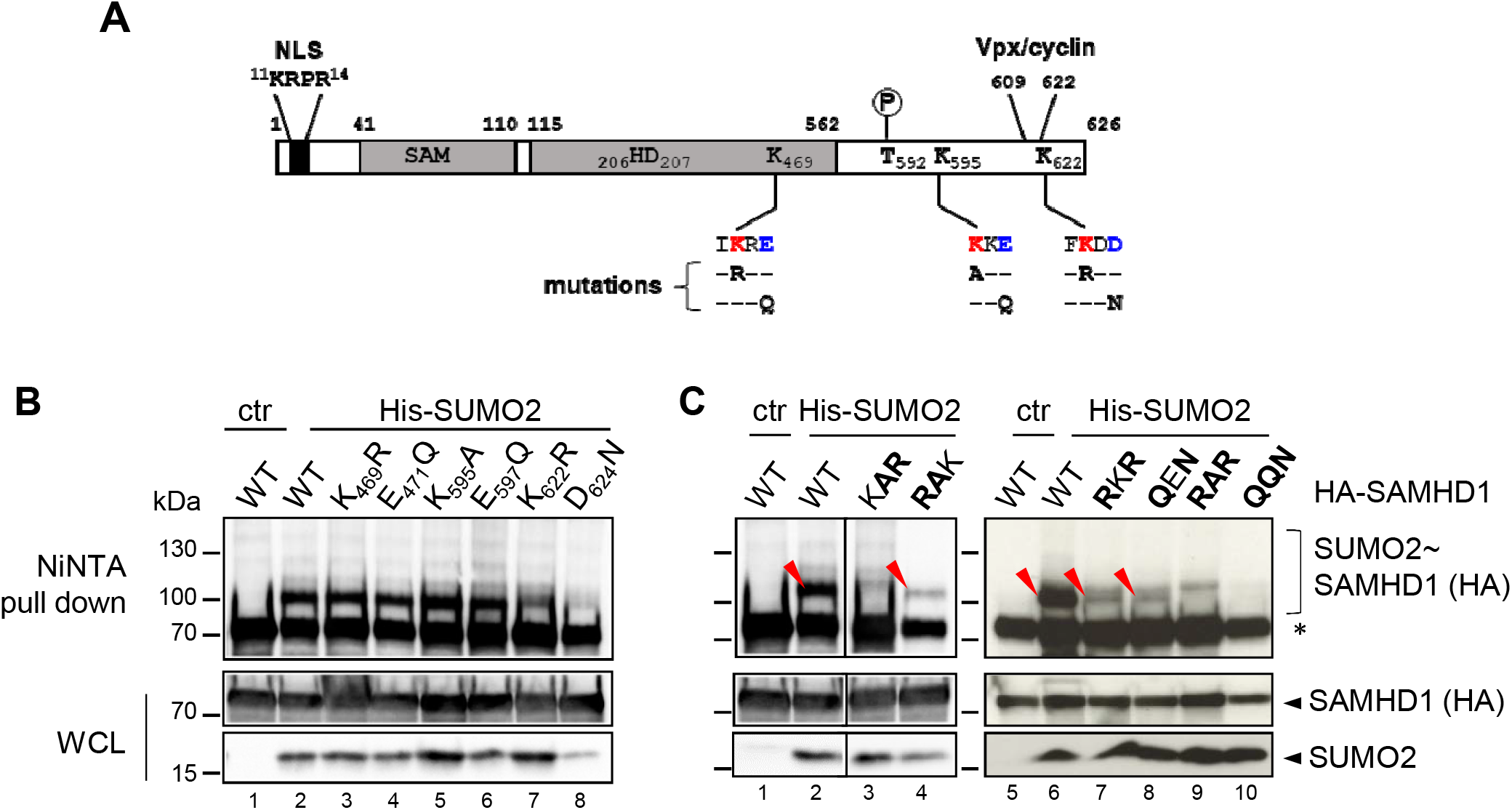
Residues K469, K595 and K622 are major SUMO conjugation sites of SAMHD1. **A.** Schematic representation of human SAMHD1 showing the nuclear localization signal (NLS), Sterile Alpha Motif (SAM) and Histidine/Aspatate (HD) domains, phosphorylatable T592 residue and the binding site for Vpx/cyclin A2. The position of the three putative SUMO consensus motifs (SCM) is indicated, with the SUMO acceptor K and the acidic amino acids colored in red and blue, respectively. Residue substitutions are described below each SUMOylation site (mutations were done on single or multiple sites as described in the text). **B.** HEK 293T cells overexpressing HA-SAMHD1 WT or single-site or **C.** multiple SUMO-site mutants, Ubc9 and His-SUMO2 were processed as in Fig. 1A. Mutated residues are in bold characters. WCL: whole cell lysate. *, nonspecific binding of unmodified SAMHD1 on Ni-NTA beads. The red arrowheads indicate the ~100 kDa band corresponding to mono-SUMOylated SAMHD1 species. Results of one representative experiment are shown (n≥2).

### SAMHD1 mutants defective for K595 SUMOylation lose their HIV-1 restriction activity

To assess the requirement of SUMO conjugation for viral restriction, we stably expressed WT or SUMOylation-site SAMHD1 mutants, in SAMHD1-negative monocytic U937 cells. Following cell differentiation by PMA treatment, all SAMHD1 mutants were enriched in the nucleus (**Fig. S5A**) and, except for the catalytic-defective variant (HD/AA), displayed WT-like expression levels (**Fig. S5B and S5C**). Next, we challenged the differentiated U937 cell lines with a VSVg-pseudotyped HIV-1 virus expressing *EGFP* as a reporter gene (VSVg/HIV-1ΔEnv*EGFP*) and quantified the fraction of infected cells 48 hours later by flow cytometry (**Fig. 3A**). As previously reported, expression of WT SAMHD1 rendered U937 cells resistant to HIV-1 infection, while the HD/AA and phosphomimetic T_592_E mutants failed to do so (**Fig. 3B**). Simultaneous substitution of the three major SUMO-acceptor K residues into R also abrogated SAMHD1-mediated restriction (**RRR** mutant, **Fig. 3B**). Similarly, preventing SUMOylation by mutation of the acidic amino acids within the corresponding SUMO consensus motifs rendered SAMHD1 restriction-defective (**QQN** mutant, **Fig. 3B**). These observations strongly indicate that the regulation of SAMHD1 antiviral activity relies on SUMOylation, but not on other K-directed PTMs (i.e. ubiquitinylation, acetylation). As SAMHD1 variants impaired for SUMO-conjugation to K469 and/or K622 efficiently blocked HIV-1 infection (**R**K**R** and **Q**E**N** mutants, **Fig. 3B** and **Fig. S5D**), we deduced that SUMOylation of K595 might be crucial for viral restriction by SAMHD1. Consistent with this hypothesis, SAMHD1 mutants where K595 was changed into either A or R lacked antiviral activity (**Fig. 3C**). Importantly, substituting E597 with Q to prevent K595 SUMOylation had similar functional consequences (**Fig. 3C**). Of note, mutating the neighboring residue Q594 into N did not modify the restriction activity of SAMHD1 (**Fig. 3C**).

**Figure 3.**
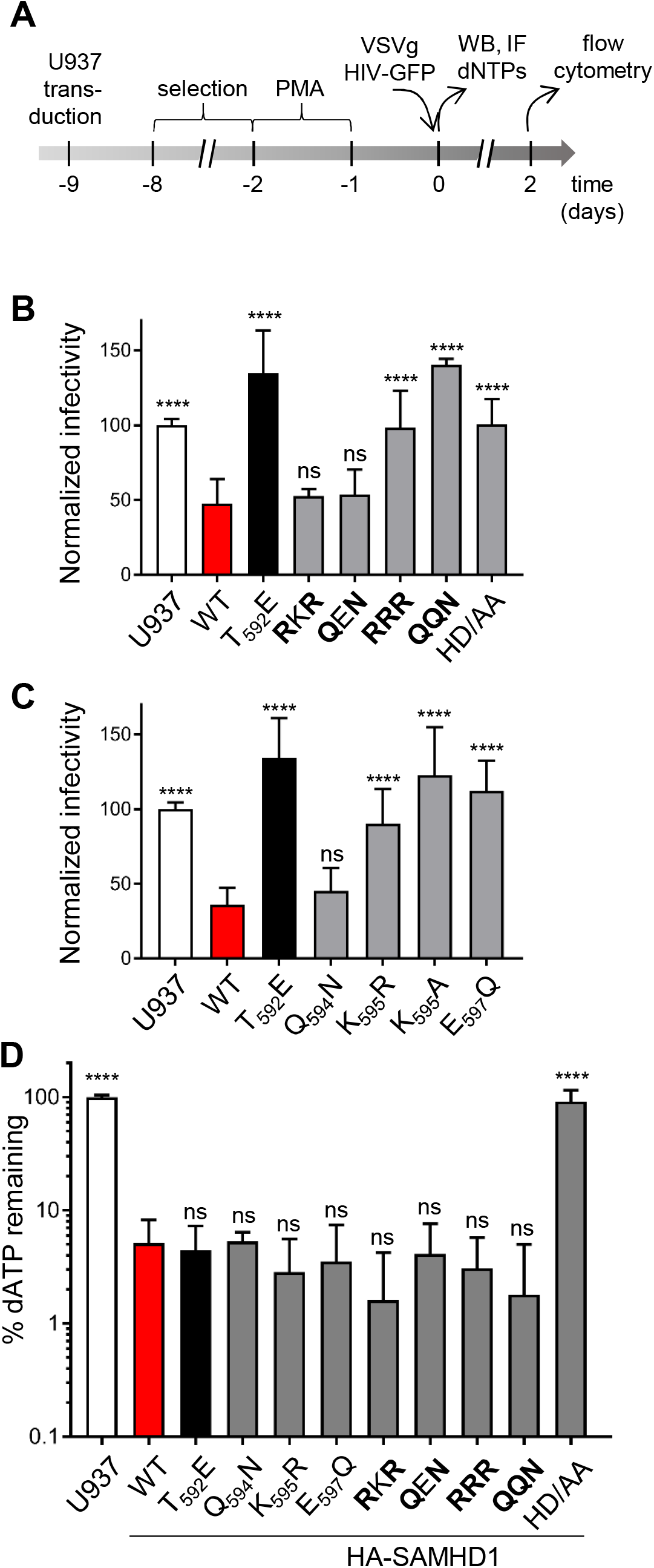
SAMHD1 mutants impaired for SUMOylation on K595 are antivirally inactive but efficiently deplete the cellular dNTP pools. **A.** U937 cell lines stably expressing HA-SAMHD1 WT or mutants (≥3 independently generated cell lines) were treated according to the experimental outline. **B.** Differentiated U937 cell lines expressing the indicated multiple or **C.** single SUMO-site SAMHD1 variants were infected with the VSVg/HIV-1ΔEnv*EGFP* virus in 3 to 4 technical replicates and analyzed by flow cytometry 48 hours later. Bold characters indicate the mutated residues as in Fig. 2. The infection rate of parental U937 cells was set to 100. Bars represent the mean ±SD (n=3 for B, 6 for C). Statistical significance was assessed by one-way ANOVA statistical test with a Dunnett’s multiple comparison post-test. ****: p<0.0001. ns: not significant. **D.** Cellular dATP levels were quantified by single nucleotide incorporation assay^9^. The dNTP levels (%) of SAMHD1-expressing cells were calculated relative to those of parental U937 set to 100. Bars show the mean ± SD (n≥3). Statistical significance was assessed by one-way ANOVA test. ****: p<0.0001. ns: not significant.

To elucidate the possible mechanisms underlying the loss-of-restriction phenotype of SAMHD1 variants defective for K595 SUMOylation, we assessed their dNTPase activity by measuring cellular dNTP levels. The concentration of dATP and dGTP (representative of the four dNTPs) of differentiated U937 cells dropped ~20-fold upon expression of WT SAMDH1 cells (**Fig. 3D** and **Fig. S5E**) reaching levels comparable to those of PMA-treated THP1 cells (**Fig. S9A**). The catalytic-defective HD/AA mutant did not alter the cellular dNTP content, while the phosphomimetic T_592_E variant was as potent as WT SAMHD1 (**Fig. 3D** and **Fig. S5E**), as previously shown^8,18,24,25^. Similarly, all the tested SUMOylation-deficient SAMHD1 mutants reduced the cellular dNTP pools to a WT extent, indicating that their dNTPase activity is intact (**Fig. 3D** and **Fig. S5E**). These results indirectly demonstrate that the impaired antiviral function of SAMHD1 mutants lacking K595 SUMOylation is due to neither defective oligomerization nor improper folding. In conclusion, impairing SUMO conjugation to K595 compromises the antiviral activity of SAMHD1 but not its dNTPase function, a phenotype that mirrors the effects of the phosphomimetic T_592_E mutation.

### Both K595 SUMOylation and viral restriction rely on the SIM2 motif

*In silico* analysis of human SAMHD1 sequence with the bioinformatic predictor JASSA^46^ highlighted the presence of three potential SIMs, suggesting that SAMHD1 could interact non-covalently with SUMO (**Fig. S6A**). SIM1 (_62_PVLL_65_) is located in the N-terminal SAM domain while SIM2 (_488_LLDV_491_) and SIM3 (spanning the overlapping _499_IVDV_501_ and _500_VDVI_502_ sequences) are found in the C-terminal half of the protein (**Fig. 4A**). Protein sequence alignment revealed that SIM1 is present in SAMHD1 orthologs from Hominids, SIM2 also in Old-World Monkey isoforms, while SIM3 is highly conserved along evolution (**Fig. S3**). By mapping the position of the putative C-terminal SIMs on the crystal structure of the HD domain of SAMHD1, we found that SIM3 is buried within the globular fold of the protein, while SIM2 is surface-exposed and near the SUMOylatable K595 residue (**Fig. 4B**), making it a more favorable candidate for functional studies. We first established that endogenous SAMHD1 was enriched on beads coupled to SUMO1 and, to a greater extent, SUMO2, but not uncoupled beads incubated with the lysate of differentiated THP1 cells (**Fig. 4C**). Next, we assessed the implication of SIM2 for the SAMHD1-SUMO binding. We found that mutating the LLDV sequence into AADA (yielding the SIM2m variant) resulted in a weaker association between SAMHD1 and SUMO2 in both pull-down (**Fig. 4D**) and PLA tests (**Fig. S6B**).

**Figure 4.**
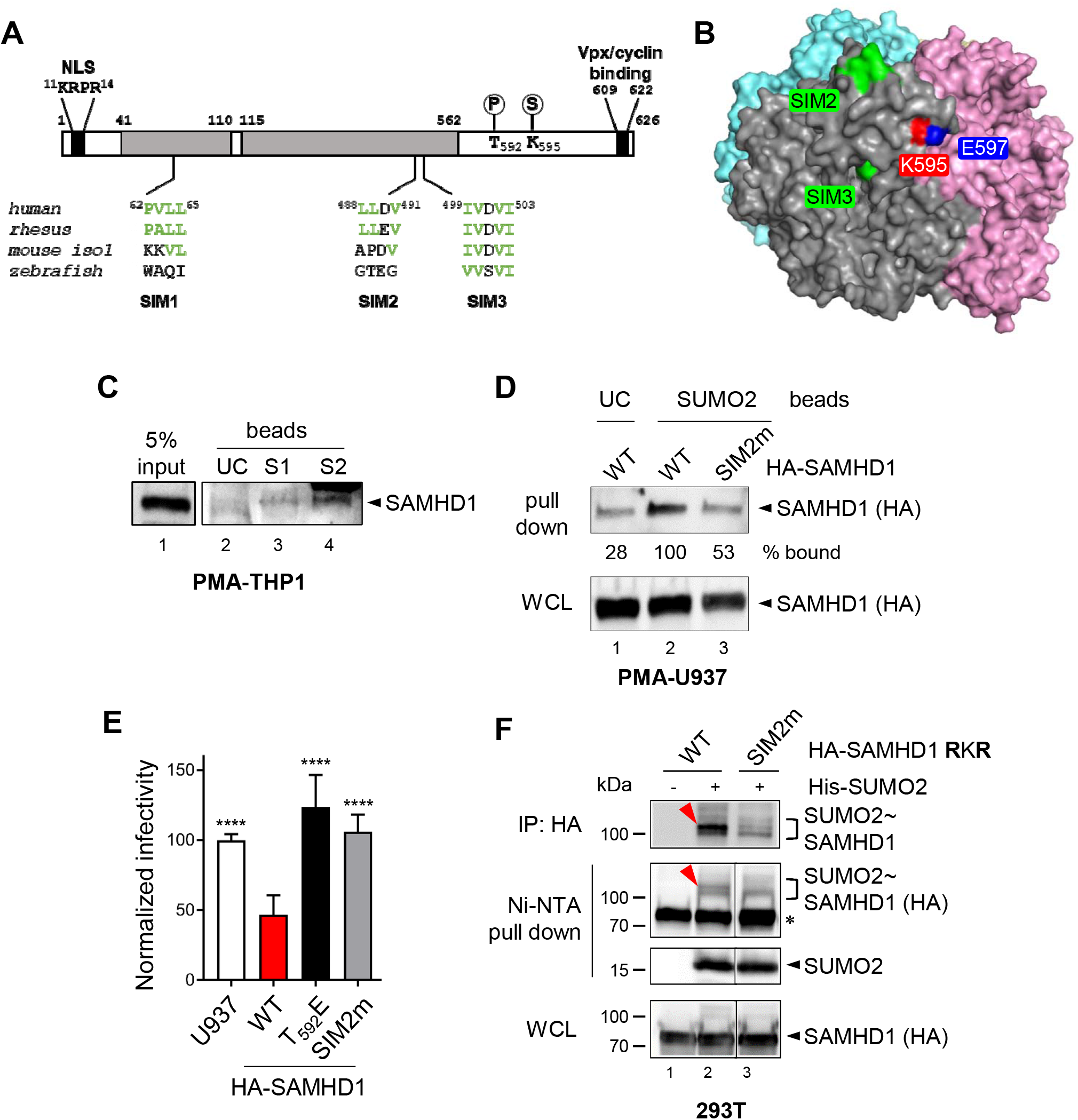
Integrity of SAMHD1 SIM2 is required for both HIV-1 restriction and K595 SUMOylation. **A.** Schematic representation of human SAMHD1 (NP_056289.2), showing the position and sequence of the putative SIMs and alignments with corresponding sequences of isoforms from Rhesus macaque (NP_001258571.1), mouse (NP_061339.3, isoform 1) and zebrafish (NP_001153405.1). **B.** Position of SIM2 and SIM3, K595 and E597 within one protomer of human SAMHD1 tetramer (PDB: 4BZC). **C.** The lysate of differentiated THP1 cells was split in equal aliquots that were incubated with agarose beads coated with either human recombinant SUMO1 (S1) or SUMO2 (S2), or uncoated beads (UC) as control. Proteins from the input and the eluates were separated by migration on a 4-15% SDS-PAGE gel and, next, visualized by immunoblotting using antibodies against SAMHD1. Results of one representative experiment are shown (n=3). **D.** The lysate of differentiated U937 cells expressing WT or SIM2m SAMHD1 variants was incubated with SUMO2-coated agarose beads and treated as in C. The band intensities were quantified with ImageJ software. Results of one representative experiment are shown (n=2). **E.** U937 cells stably expressing the indicated HA-SAMHD1 variants (3 independently generated cell lines) were infected with the VSVg/HIV-1ΔEnv*EGFP* virus in 4 technical replicates and analyzed as in figure 3B. Bars represent the mean ±SD (n=4). The infection rate of parental U937 cells was set to 100. Statistical significance was determined by one-way ANOVA test. ****: p<0.0001. **F.** HEK 293T cells overexpressing HA-SAMHD1 **R**K**R** mutant, Ubc9 and His-SUMO2 were lyzed in denaturing conditions and split in two equal aliquots that were subject to affinity purification on either Ni-NTA or HA-matrix beads. Eluted proteins were analyzed as described in Fig. 1A. WCL: whole cell lysate. Results of one representative experiment are shown (n=2). The red arrowhead highlights the SUMO-conjugated K595 SAMHD1 specie. *, nonspecific binding of unmodified SAMHD1 on Ni-NTA beads.

The existence of a non-covalent interaction between SAMHD1 and SUMO mediated by SIM2 prompted us to investigate the possible implications for restriction using the U937 cell-based assay described above. Mutation of SIM2 rendered SAMHD1 unable to inhibit both HIV-1 (**Fig. 4E**) and HIV-2ΔVpx infection (**Fig. S7A**) without affecting its localization (**Fig. S6B**), expression levels (**Fig. S7B**) and capacity to reduce cellular dNTP concentrations (**Fig. S7C**). SIM2 is located near residue T592, which phosphorylation is recognized as a major regulator of SAMHD1 antiviral function^17,18^. Therefore, we used an anti-phospho-T592 species-specific antibody to monitor the degree of modification of SAMHD1 SIM2m mutant expressed either stably in differentiated U937 cells or transiently in cycling 293T cells. In both cell types, SAMHD1 SIM2m variant was phosphorylated at levels comparable to that of WT SAMHD1 (**Fig. S7B and S7D)**.

As mutation of either SIM2 or the adjacent SUMOylation motif harboring K595 abrogated SAMHD1 restriction activity, we postulated the existence of a functional connection between the two sites. Indeed, SIM2 might provide an extended binding interface that stabilizes the association between SAMHD1 and the SUMO-charged Ubc9 promoting the efficient transfer of SUMO to K595, which lies within a minimal SUMOylation site (KxE)^34^. By performing both immunoprecipitation and histidine affinity purification assays, we confirmed that the LLDV/AADA substitution virtually abolished SUMO conjugation to SAMHD1 **R**K**R** variant, as demonstrated by the loss of the ~100 kDa band corresponding to K595 SUMOylation (**Fig. 4F**, compare lanes 2 and 3). Conversely, inactivation of SIM2 barely modified the conjugation profile of WT SAMHD1 in conditions of either ectopic (**Fig. S7E**, compare lanes 2 and 3) or endogenous expression of SUMO isoforms (**Fig. 1B**, compare lanes 2 and 3). In conclusion, the integrity of the surface-exposed SIM2 is essential for both SUMO conjugation to K595 and viral restriction, providing converging evidence that human SAMHD1 requires K595 SUMOylation to be restriction competent.

### Modification of K595 by SUMO2 renders dephosphorylated SAMHD1 antivirally active

SAMHD1 mutants defective for SUMO conjugation to K595 mirrored the loss-of-restriction phenotype of the phosphomimetic T_592_E variant, suggesting that SUMOylation and phosphorylation of these adjacent sites might interfere with each other. We reasoned that T592 phosphorylation might inhibit modification of K595 by SUMO either directly, by introducing a negative charge that generates an electrostatic repulsion, or indirectly, by modifying the conformation of the C-terminal region of SAMHD1 encompassing residues 582-626^23,47^. The SUMOylation profile of the SAMHD1 **R**K**R** mutant was unaffected if T592 was changed into A or E, to mimic the absence or presence of a phosphate group, respectively (**Fig. S8A**). Therefore, the phosphorylation status of T592 does not seem to affect K595 SUMOylation.

We then analyzed the degree of phosphorylation of WT and SUMOylation-deficient SAMHD1 variants expressed in 293T cells using a phospho-T592 species-specific antibody. The E_597_Q change did not detectably modify the ratio of the phosphorylated relative to the total SAMHD1 levels, indicating that K595 SUMOylation is not a requirement for T592 phosphorylation. We also observed that the Q_594_N and K_595_R substitution reduced T592 phosphorylation by ~33 to 47%, respectively (**Fig. 5A**). Strikingly, the K_595_A mutation, which rendered SAMHD1 antivirally inactive (**Fig. 3C**), caused a ~94% drop in T592 phosphorylation (**Fig. 5A**), likely due to the inactivation of the ^592^TPQK^595^ CDK-consensus sequence. We obtained similar results in differentiated U937 cells (**Fig. S8B**). Lack of T592 phosphorylation upon replacing K595 with alanine was confirmed by comparing the electrophoretic mobility of WT and SUMOylation-deficient SAMHD1 variants in a Phos-tag gel, ruling out the possibility that amino acids changes near T592 had altered the binding affinity of the phospho-specific antibody (**Fig. 5B**). The absence of correlation between the status of phosphorylation and SUMOylation of SAMHD1 suggest that these PTMs are independent of one another. Importantly, these observations show for the first time that dephosphorylated T592 alone is insufficient to render SAMHD1 restriction competent. To substantiate further this conclusion, we generated U937 cell lines stably expressing SAMHD1 variants bearing the phospho-ablative T_592_A change alone or together with mutations disrupting SUMO conjugation to K595 (**Fig. S8C**) and tested their restriction activity. As previously reported^17,18,23,48^, SAMHD1 T_592_A mutant restricted HIV-1 to the same extent as its WT counterpart (**Fig. 5C**). Concomitant mutation of K595 rescued HIV-1 infectivity to a moderate but statistically significant extent (**Fig. 5C**), a phenotype that might be attributed to the transfer of SUMO to an adjacent site (i.e. K596). The E_597_Q change had a stronger effect, turning the constitutive-active SAMHD1 T_592_A variant into a restriction-defective protein (**Fig. 5C**).

**Figure 5.**
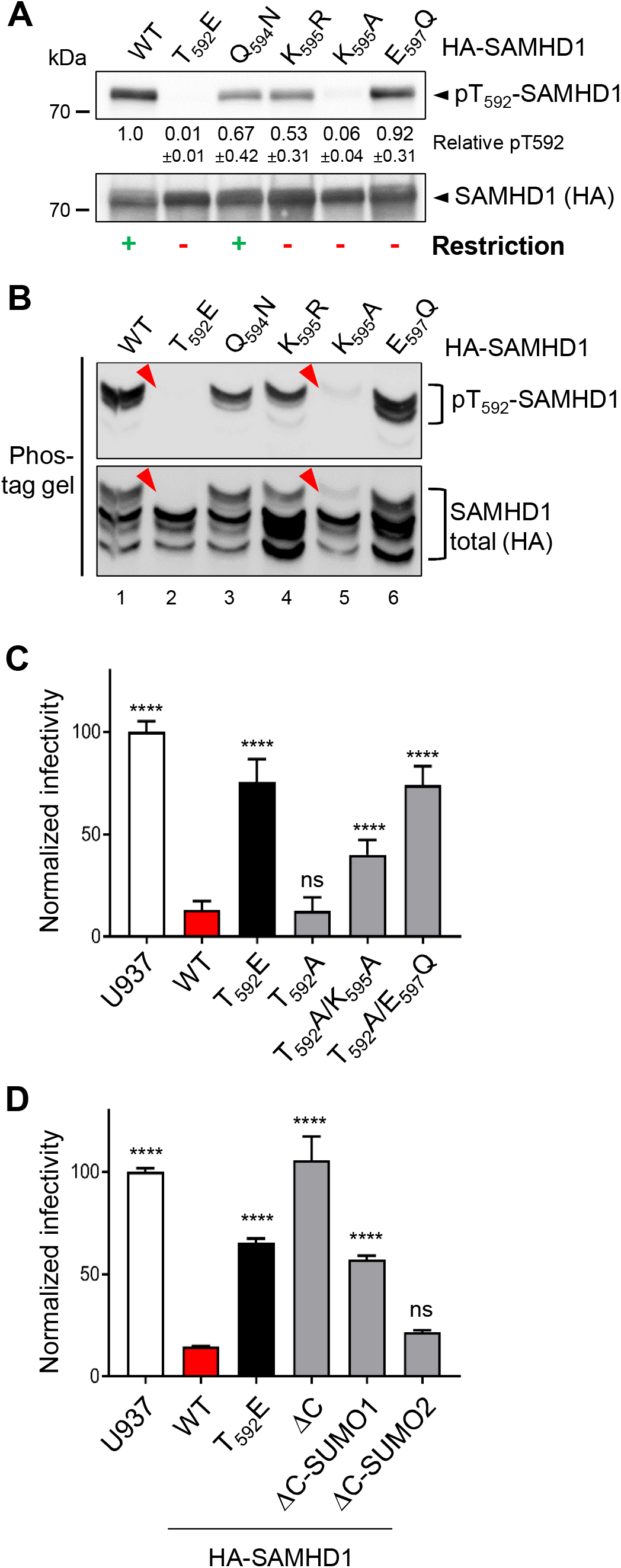
Dephosphorylated T592 is not sufficient to render SAMHD1 antivirally active, concomitant SUMOylation of K595 is required. **A.** Proteins (10 μg total proteins/line) contained in the crude extract of 293T cells overexpressing HA-SAMHD1 variants were loaded on a 4-15% pre-casted SDS-PAGE gel. Immunoblotting was performed sequentially with an anti-pT592 or anti-HA antibody to detect either phosphorylated or total SAMHD1 species, respectively. The band intensities were quantified by densitometry with ImageJ software and the pT592/total SAMHD1 ratio for WT SAMHD1-expressing cells was set to 1. Results of one representative experiment are shown (n=2). The ability of SAMHD1 mutants to restrict (green +) or not (red -) viral infection is indicated. **B.** The same samples as in A were separated on a 7% Phos-tag^™^ SDS-PAGE gel. Arrowheads indicate that T592 phosphorylated SAMHD1 species which become undetectable upon T592E or K595A mutation. Results of one representative experiment are shown (n=3) **C.** and **D.** U937 cells stably expressing the indicated HA-SAMHD1 mutants were infected with the VSVg/HIV-1ΔEnv*EGFP* virus in 4 technical replicates and analyzed by flow cytometry 24 hours later. The infection rate of parental U937 cells was set to 100. Bars represent the mean ±SD (n=2). Statistical significance was determined by one-way ANOVA test. ****: p<0.0001. ns: not significant.

Previous studies showed that the deletion of the C-terminal tail (aa 595-626, SAMHD1 ΔC) renders SAMHD1 restriction-defective^23,49^. To exclude that mutation of T592 together with K595 or E597 might had altered this functionally important region of SAMHD1, we engineered two fusion proteins where either the SUMO1 or the SUMO2 sequence was inserted in-frame at the C-terminus of the SAMHD1 ΔC truncation mutant to mimic a constitutively SUMOylated form. We reasoned that, if the defect was due to lack of K595 SUMOylation, the SUMO-fusion should compensate for the loss of function of the SAMHD1ΔC mutant, as reported for CtIP-dependent DNA resection activity^50^. Using the U937-cell based restriction assay we confirmed that SAMHD1ΔC was unable to inhibit HIV-1 infection (**Fig. 5D**). Importantly, fusion of SUMO2, but not SUMO1, to SAMHD1ΔC fully restored viral restriction (**Fig. 5D**), with no detectable change of the protein expression levels (**Fig. S8D**). Collectively, our data demonstrate that, in non-cycling cells where the bulk of SAMHD1 harbors dephosphorylated T592, SUMOylation of K595 is required for viral restriction.

### Infectivity of SAMHD1-sensitive viruses raises upon SUMOylation inhibition in macrophages

Having shown that SUMOylation of the conserved K595 residue regulates the antiviral activity of SAMHD1 stably expressed in U937 cells, we undertook a drug-based approach to validate this finding on endogenous SAMHD1-expressing cells. We therefore exposed undifferentiated or PMA-treated THP1 cells to GA or AA, to suppress the activity of the E1 SUMO-activating enzyme^42^ and hinder the SAMHD1-SUMO association (**Fig 1D**), before challenge with the VSVg/HIV-1ΔEnv*EGFP* virus. Cycling THP1 cells were readily permissive to HIV-1 and inhibition of SUMOylation did not alter this state (**Fig. 6A**), indirectly demonstrating that cell viability was not affected. The percentage of HIV-1-positive cells sharply decreased (~15-fold) following differentiation and was rescued to a moderate (~1.7 to 2-fold) but statistically significant extent by blocking the SUMO pathway (**Fig. 6A**). This effect was neither due to altered SAMHD1 localization (**Fig. 1D**) and expression (**Fig. S1**), nor changes in the dNTP concentrations (**Fig. S9A**). The implication of the SUMOylation process in the antiviral mechanism of SAMHD1 was further substantiated by finding that treatment with GA increased ~2.5-fold the permissiveness of differentiated THP1 cells to HIV-2ΔVpx without affecting the infectivity of the corresponding WT virus (**Fig. 6B**).

**Figure 6.**
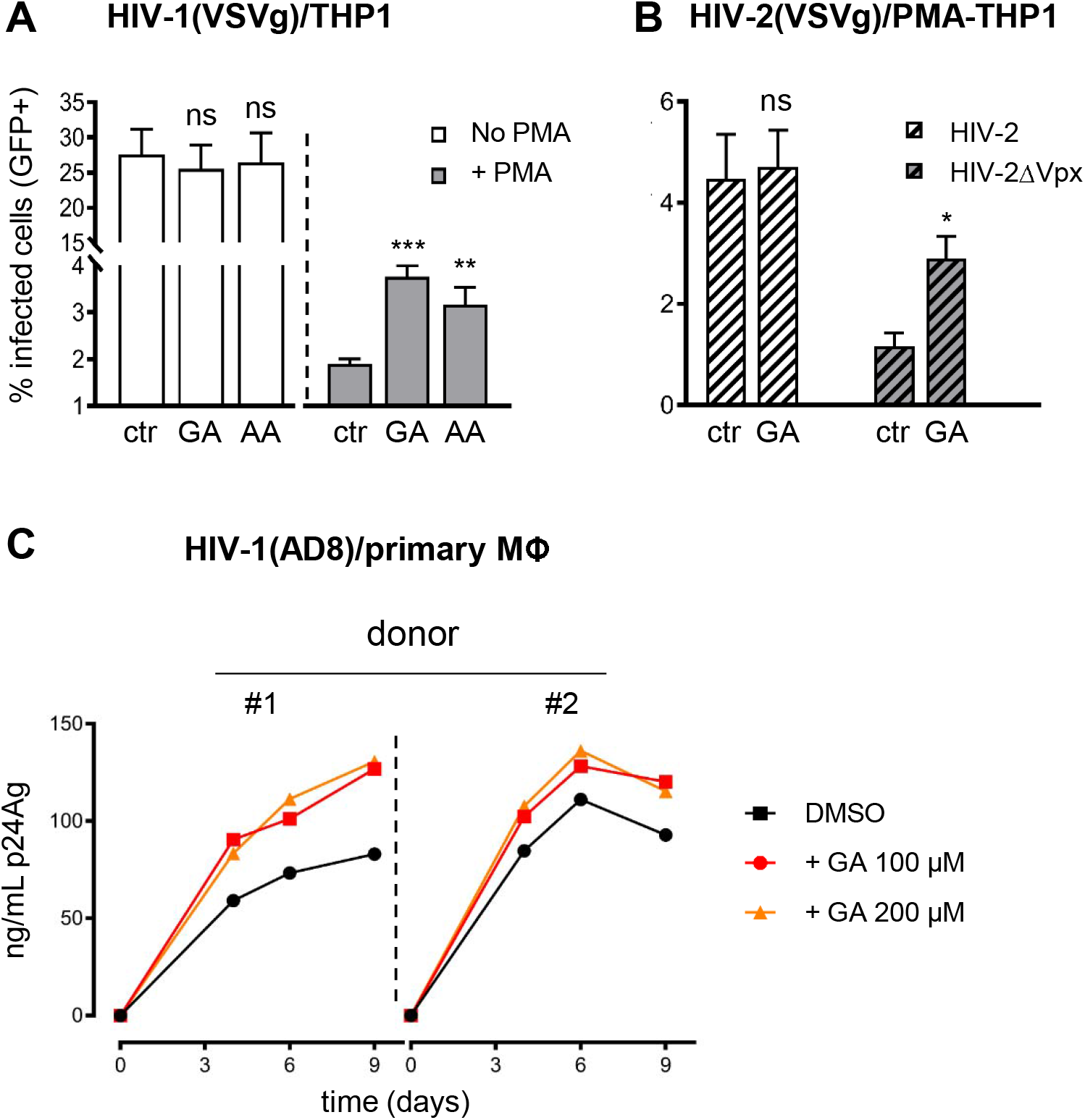
Infectivity of SAMHD1-sensitive viruses is enhanced by inhibition of SUMOylation in macrophages. **A.** THP1 cells were differentiated into macrophage-like cells by incubation with PMA (100 ng/mL, 24 h), or left untreated. Twenty-four hours later SUMOylation was inhibited by exposure to gingkolic acid (GA, 50μM) or anacardic acid (AA, 50μM) for 2 hours before challenge with VSVg-pseudotyped HIV-1 (moi = 0.2), **B.** HIV-2 or HIV-2ΔVpx (moi = 0.3) viruses harboring the *EGFP* reporter gene. The percentage of infected (GFP-positive) cells was measured by flow cytometry after 48 hours. Bars represent mean ± SD of 3 technical replicates. One representative experiment is shown (n=5 for A, 2 for B). Statistical significance was assessed by 2 way-ANOVA with Sidak’s multiple comparisons test. *: p<0.05; **: p<0.01; ***: p<0.001; ns: not significant. **C.** Monocyte-derived macrophages generated from 2 healthy donors were pretreated with GA or vehicle (DMSO, 2h) before challenge with the AD8 HIV-1 strain (10 ng/mL p24Ag). Viral replication was monitored overtime by measuring the p24 antigen (p24Ag) released in the cell culture supernatant.

Finally, we investigated the outcome of GA treatment on the spread of the replication-competent HIV-1 AD8 macrophage-tropic virus in primary monocyte-derived macrophages (MDMs). Inhibition of SUMOylation enhanced the magnitude of HIV-1 viral particle release in cultures of MDMs from 3 different donors, although to a various degree (**Fig. 6C** and **S9B**). Similar results were recapitulated upon infection of induced-pluripotent stem cells (iPSc)-derived macrophages (**Fig. S9C**). Altogether, these observations confirm that the maintenance of a SAMHD1-dependent antiviral state in macrophages relies on a functional SUMO system.

## Discussion

It is widely accepted that the antiviral activity of SAMHD1 is downregulated by phosphorylation of residue T592 in actively dividing cells. Yet, T592 phosphorylation does not influence the dNTPase function, which is a central element of the restriction mechanism mediated by SAMHD1, implying that additional activities and/or regulations are at play. In this study, we show that SAMHD1 is SUMOylated and provide several lines of evidence that this modification critically regulates its antiviral activity in non-cycling immune cells. Indeed, we demonstrate that SAMHD1 is conjugated by the three SUMO paralogs, expressed either ectopically or at endogenous levels, in cycling cells. Combining biochemical and imaging approaches, we also show that modification of SAMHD1 by SUMOs occurs in the nucleus of differentiated cells of myeloid origin, where its antiviral function is witnessed. Next, we identify residues K469, K595 and K622 as the major SUMO-attachment sites of SAMHD1. Still, we cannot exclude that additional sites might be SUMOylated at a low level or under specific circumstances (**Table S1**). By comparing the SUMO modification profile of SAMHD1 variants, where these amino acids were mutated in various combinations, we found that K595 and K622 undergo mono-SUMOylation, while K469 and K622 are targeted by SUMO chains, which accumulate upon inhibition of the proteasome (**Fig. 7A**). Recently, ubiquitination of K622 mediated by TRIM21, which belongs to a protein family comprising ligases with dual SUMO and Ubiquitin E3 activity^51^, was proposed to promote proteasomal degradation of SAMHD1 in enterovirus-infected cells^52^. Whether Ubiquitination and SUMOylation of the same site cooperate to regulate the fate of SAMHD1 remains open for future studies. We also discovered that SUMOylation of K595, which is embedded within a minimal SUMO-consensus motif (KxE), relies on the integrity of a proximal SIM (named SIM2, aa 488-491). The presence of a SIM might contribute to an efficient recruitment of the SUMO-charged Ubc9, thereby promoting SUMOylation of the adjacent K595 residue^34^. By mediating a preferential non-covalent interaction between SAMHD1 and SUMO2 (**Fig. 4C**), SIM2 might also dictate the selective modification of K595 by this paralog which is needed for viral restriction (**Fig. 5D**).

**Figure 7.**
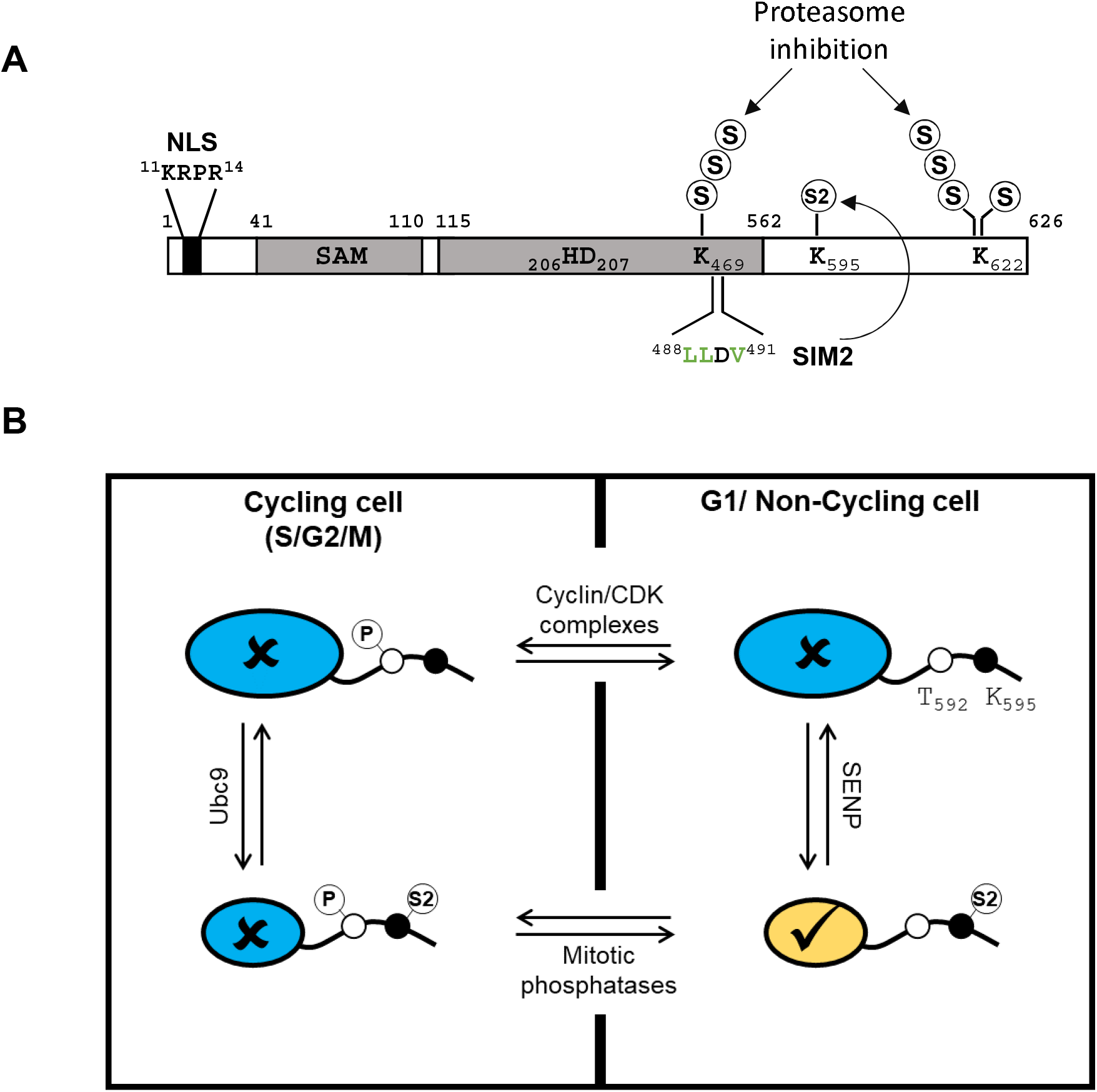
Model for the regulation of SAMHD1 antiviral activity by SUMOylation in nondividing cells. **A.** Human SAMHD1 harbors three major SUMO-attachment sites: K595 and K622 undergo mono-SUMOylation, while K469 and K622 are targeted by SUMO chains, which accumulate upon inhibition of the proteasome. Human SAMHD1 also harbors the surface-exposed SIM2 sequence, which drives the modification of K595 by SUMO, likely with a preference for the SUMO2 isoform (S2). **B.** In actively dividing cells, SAMHD1 is targeted by CDK/cyclin-mediated phosphorylation on T592 during the G1/S transition thereby losing its antiviral activity (✗, colored in blue). A fraction of SAMHD1 (smaller circle) might be SUMOylated on K595 by the action of Ubc9. However, this modification appears insufficient to neutralize the effects of phosphorylation and rescue restriction. Upon mitotic exit, phosphorylation is reversed by host PPP family phosphatases. Our results show that SAMHD1 harboring dephosphorylated T592 is antivirally inactive if SUMO-conjugation to K595 is prevented. They also indicate that only the fraction of SAMHD1 that harbors SUMOylated K595 and dephosphorylated T592 (✔, colored in yellow) inhibits viral infection through a dNTPase-independent mechanism.

By interrogating the ability of SUMOylation-deficient SAMHD1 variants to inhibit HIV-1, we found that simultaneous mutation of the three SUMO-acceptor K residues abolishes viral restriction. The same was observed upon substitution of the acidic amino acids within the corresponding SUMO-consensus motifs, to prevent the recruitment of the SUMOylation machinery^36,37^, strongly supporting the implication of SUMO conjugation, but not other K-directed PTMs, for the antiviral function. Further investigations showed that inactivation of the SUMO-consensus site harboring K595 alone recapitulates the loss-of-restriction phenotype. Similarly, mutation of SIM2, which hampers SUMOylation of K595 (**Fig. 4F**), renders SAMHD1 unable to restrict both HIV-1 and HIV-2ΔVpx. These results provide converging evidence that viral restriction relies on SUMOylation of K595, while modification of K469 and K622 by SUMO is dispensable.

In a complementary approach, we assessed whether small molecule SUMOylation inhibitors, which weakened the SAMHD1-SUMO interaction (**Fig. 1D**), might relieve the restriction activity of endogenous SAMHD1 and promote infection. Blocking SUMOylation enhanced the cell permissiveness to infection in contexts where SAMHD1 was antivirally active (differentiated THP1 cells infected by either HIV-1 or HIV-2ΔVpx), but not when its function was suppressed by either T592 phosphorylation (dividing THP1 cells, HIV-1 infection) or Vpx-mediated degradation (differentiated THP1 cells, infection by HIV-2). Importantly, inhibiting SUMOylation favored the spreading of a replication-competent HIV-1 virus in cultures of primary human MDMs indicating, on the side, that the observed effects are independent on the viral entry pathway. These results support the implication of SUMOylation in the antiviral response mediated by SAMDH1.

Several data indicate that the restriction mechanism of SAMHD1 is tightly connected to its ability to limit the dNTP supply for viral genome replication^4,8,23^. However, all the SUMOylation-defective mutants tested in this study (including K_595_A, K_595_R, E_597_Q and SIM2m) were as potent as WT SAMHD1 in reducing the cellular dNTP concentrations. Along this line, SUMOylation inhibitors increased the permissiveness of macrophages to SAMHD1-sensitive viruses without altering the cellular dNTP levels. We concluded that SUMO conjugation does not influence the ability of SAMHD1 to hydrolyze dNTPs in cells. The possibility to uncouple the modulation of the dNTP pools and the antiviral function, which was previously described for the phosphomimetic T_592_E^17,18,24,25^ and the redox-insensitive C_522_S^53^ variants, strongly indicates that SAMHD1 possesses additional dNTPase-independent properties contributing to the viral restriction mechanism, which await identification.

The covalent attachment of SUMO to K595 might directly stimulate another enzymatic function of SAMHD1 relevant for virus restriction, i.e. the debated RNAse activity^24,28,29,54^. Nevertheless, SUMOylation generally occurs at a low stoichiometry (often less than 1%^55^), which makes it difficult to envision how preventing the modification of a tiny proportion of SAMHD1 could account for the dramatic restriction defect that we observed. Notably, the ability of SAMHD1 to interact with the viral genome^27^ could favor SUMOylation, as reported for PCNA^56^ and PARP-1^57^, and influence, in turn, its SUMO-dependent antiviral functions. Another major consequence of SUMOylation is the formation of new binding interfaces, leading to the notion that SUMO acts as a “molecular glue” between its substrate and a SIM-containing partner, which otherwise display weak affinity^58^. Thus, we speculate that SUMOylation of K595 might stimulate the recruitment of cellular and/or viral cofactors via their SIM, bringing about the formation of a complex endowed with antiviral activity. Phosphorylation of T592 might interfere with the assembly of such a complex. In an alternative scenario, SUMO as a bulky modifier might abrogate the association between SAMHD1 and an inhibitor keeping it in an antivirally inactive state. Further investigation is warranted to investigate these hypotheses.

The analogous restriction phenotype of SAMHD1 variants harboring mutations that either mimic a constitutively phosphorylated T592 or impair SUMO conjugation to K595 raised the possibility that these PTMs might be co-regulated. However, the lack of correlation between the phosphorylation and the SUMOylation status of SAMHD1 suggests that these PTMs are independent events. On one hand, this implies that phosphorylation of T592, which is important for replication fork progression and resection of collapsed forks^21^, abolishes the restriction activity of SAMHD1 in cycling cells, irrespective of whether K595 is SUMOylated or not (**Fig. 7B**). On the other hand, our results support a model according to which K595 SUMOylation defines the subpopulation of restriction-competent SAMHD1 in non-cycling immune cells, where the bulk of the protein harbors a dephosphorylated T592 residue. First, we found that SAMHD1 K_595_A mutant displayed a severe phosphorylation deficiency, possibly because the CDK-consensus driving T592 phosphorylation is disrupted^43^. As SAMHD1 K_595_A variant lacked restriction activity, these data indicate that dephosphorylated T592 alone cannot overcome the defect imposed by the absence of K595 SUMOylation (**Fig. 7B**). Second, we confirmed that the K_595_R or E_597_Q change, which inhibit SUMO attachment to K595, converted the constitutively active SAMHD1 T_592_A variant into a restriction-defective protein. Third, we found that the restriction activity of the SAMHD1 ΔC mutant (lacking aa 595-626) was fully rescued upon fusion with SUMO2. Conversely, SUMO1 had only a mild effect, despite ~50% sequence homology with its paralog^35^. These isoform-specific effects are intriguing since SAMHD1 displays a preferential non-covalent interaction with SUMO2 rather than SUMO1 (**Fig. 4C**). Moreover, it should be noted the fraction of SUMO2/3 available for conjugation is larger than that of SUMO1, which is mostly conjugated to high-affinity targets (i.e. RanGAP1), and can further raise in response to various stimuli including viral and bacterial infection^33,44,59^.

In conclusion, our results unravel that the regulation of the antiviral function of SAMHD1 depending on the cell cycle status of the infected cell is more complex than previously anticipated and point to a scenario where phosphorylation of T592 and SUMOylation of K595 provide a sophisticated mechanism controlling a dNTPase-independent component of the restriction activity. These findings not only open a new perspective to uncover the enigmatic aspects of SAMHD1-mediated viral restriction, but also provide opportunities for the development of strategies aiming to selectively manipulate the immune function of SAMHD1 without affecting activities important for cell homeostasis.

## Methods

### Cells and reagents

Human Embryonic Kidney (HEK) 293T cells were cultured in DMEM (Invitrogen). The human monocytic U937 and THP1 cell lines were grown in RPMI (Invitrogen). Media were supplemented with 10% fetal calf serum (Invitrogen) and penicillin/streptomycin (100 U/mL). U937 and THP1 cell lines were differentiated by treatment with phorbol-12-myristate-13-acetate (PMA, Sigma-Aldrich) (100 to 300 ng/mL, 24h). All cell lines were tested mycoplasma-free (Mycoplasmacheck, GATC Biotech). Buffy coats from human healthy donors were obtained from the “Etablissement Français du Sang”. Monocytes were isolated using a CD14+ selection kit (Miltenyi Biotech) and cultured 12 days in DMEM supplemented with 10% Human Serum (inactivated) to generate MDMs. Antibodies used are the following: mouse anti-HA11 (Covance), sheep anti-SUMO1 (Enzo), rabbit anti-SUMO1, rabbit anti-SUMO2/3, mouse anti-SAMHD1, anti-HA HRP (Abcam), rabbit anti-pT592-SAMHD1 (Cell Signaling), rabbit anti-actin, anti-Flag HRP (Sigma-Aldrich), anti-HA HRP (Roche).

### Plasmid construction and mutagenesis

pMD2.G encodes the VSVg envelope protein and psPAX2 is a second-generation HIV-1-based packaging plasmid (a gift from D. Trono). pNL4-3EnvFsGFP contains a complete HIV-1 provirus with an *env*-inactivating mutation and *EGFP* inserted in the place of the Nef-coding gene (a gift from D. Gabuzda)^60^. HIV-2 ROD9Δenv-GFP (WT or ΔVpx) was described previously^61^. HIV^NLAD8^ is a macrophage (CCR5) tropic HIV-1 derivative of pNL4-3 containing the ADA envelope^62^. His-SUMO1, 2 and 3 were already described^63^. pLenti-puro construct expressing N-terminal HA-tagged human SAMHD1 was described previously^8^ and was used as template to generate mutants using the Q5^®^ Site-Directed Mutagenesis Kit according to the manufacturer’s instructions (NEB). To obtain the SAMHD1ΔC-SUMO fusion proteins, a SalI site was inserted at position 1791 in the coding sequence of SAMHD1 (between K596 and E597). The ORF encoding SUMO1 or SUMO2 flanked by XhoI and SalI sites was amplified by PCR and then inserted by ligation into the modified vector digested by SalI. The C-terminal GG motif of SUMOs was mutated into AA to prevent conjugation. The entire coding fragment was confirmed by sequencing (GATC Biotech).

### Virus stock production, infection assay, stable cell lines

Single-round viruses were produced by co-transfection of 293T cells using a standard calcium phosphate precipitation technique with the pNL4-3EnvFsGFP or HIV-2 ROD9ΔenvGFP plasmids and a VSVg-expression vector (pMD2.G) at a 20:1 ratio. Supernatants were collected 48h post-transfection, clarified by centrifugation, filtered through 45 μm-pore size filters and concentrated onto a 20% sucrose cushion by ultracentrifugation (24,000 rpm, 2h, 6°C) using a SW32 rotor (Beckman). HIV-1-based lentiviral particles were produced by co-transfecting 293T cells with packaging (psPAX2), VSVg-expressing (pMD2.G) and vector plasmids at a 4:1:5 ratio. Stable U937 cell lines were subject to puromycin selection (4 μg/mL, 6 days). Infection assays were conducted in a 12-well plate (0.3×10^6^ cells/well) in 3 to 4 technical replicates and the percentage of GFP-expressing cells was quantified after 24 or 48 hours on a FORTESSA flow cytometer using a BD FACS DIVA software (BD Biosciences). Inocula were adjusted to yield ~30% GFP-positive PMA-treated U937 cells, corresponding to a theoretical multiplicity of infection (moi) of 0.3. MDMs were seeded in flat-bottomed 96-well plates at 1×10^6^ cells per well. Following incubation with GA (2h) cells were infected with HIV^NLAD8^ (10ng/mL), and the viral p24 antigen released in the supernatant was quantified at day 4, 6 and 9 post-infection.

### Denaturing purification on Ni-NTA beads and immunoprecipitation assays

293T cells (3×10^6^ cells/10-cm dish) were transfected using a calcium phosphate precipitation technique with plasmids encoding HA-tagged WT or mutant SAMHD1 proteins, Ubc9 and each SUMO paralog bearing an N-terminal 6-His tag or an appropriate empty plasmid. When required, cells were treated with MG132 overnight (ON, 3 μM, Merck). Cells were lysed in ice-cold RIPA buffer (150mM NaCl, 0,4% NaDOC, 1% IGEPAL CA-630, 50mM Tris HCl pH 7.5, 5mM EDTA, 10mM NEM, 1mM DTT, proteases cocktail inhibitors) supplemented with 1% SDS and 1% TritonX-100. Following dilution 1:5 in RIPA buffer, lysates were incubated on HA-Tag affinity matrix beads (Pierce^™^) (ON, RT). Alternatively, cells were lysed in buffer A (6M guanidium-HCl, 0.1M Na_2_HPO_4_/NaH_2_PO_4_, 10mM imidazole, pH 8.0) and sonicated with a Bioruptor^™^ (Diagenode) (10 cycles, 45” pulse, 20” pause) before incubation with Nickel-Nitrilotriacetic (Ni-NTA) agarose beads (QIAGEN) (3h, RT). Following extensive washing with decreasing concentrations of guanidium-HCl, bound proteins were eluted by boiling in Laemmli buffer supplemented with 200 mM imidazole. For IP assays in native conditions, cells were lysed in ice-cold buffer (150 mM Tris HCl pH 8.0, 150 mM NaCl, 1% Triton, proteases cocktail inhibitors) and sonicated. Pre-cleared cell lysates were incubated with agarose beads coupled with either Human recombinant SUMO-1 or SUMO-2 (Enzo) (ON, 4°C). Following extensive washing in lysis buffer, bound proteins were eluted by boiling in Laemmli buffer.

### Total protein quantification and immunoblotting

The total protein concentration was determined by Lowry’s method using the DC Protein Assay Kit, according to the manufacturer’s instructions (Bio-Rad) with serial dilution series of Bovine Serum Albumin (BSA, Sigma) used as calibration standard. The optical density was measured at 750 nm using a plate reader (Berthold) with MikroWin 2010 software. Proteins contained in the whole cell lysate (WCL) or the eluates were separated on a 4-15% gradient sodium dodecylsulfate-polyacrylamide gel electrophoresis (SDS-PAGE) pre-casted gel (Bio-Rad). For Mn^2+^-Phos-tag SDS-PAGE gel, the acrylamide-pendant Phos-tag ligand (50 μM, Phos-tag^™^ Acrylamide, FUJIFILM Wako Chemicals USA Corporation) and MnCl_2_ (100 mM) were added to the 7% separating gel before polymerization. The Phos-tag gel was soaked in a transfer buffer containing 5 mM EDTA (10 min x 3) followed by washing in a transfer buffer without EDTA (20 min). Proteins were transferred on a nitrocellulose membrane and then detected with appropriated antibodies. Immune-complexes were revealed with HRP-conjugated secondary antibodies and enhanced chemoluminescence (Pierce^™^ ECL Western Blotting Substrate).

### Immunofluorescence, in situ proximity ligation assays (Duolink), and confocal microscopy

Cells seeded into 8-chamber culture slides (Nunc^™^ Lab-Tek^™^ II Chamber Slide^™^ System, ThermoFisher Scientific) (0.4×10^6^ cells/well) were fixed (4% paraformaldehyde/PBS, 15’, 4°C), permeabilized (0.1% PBS-Triton, 20’, 4°C), quenched (125 mM glycin) and incubated with primary antibodies (ON, 4°C) diluted in blocking buffer (5% BSA, 0.1% PBS-Tween20), followed by secondary antibody coupled to Alexa_594_ or Alexa_488_ dye (1h, RT). Protein-protein interactions *in situ* were visualized using the Duolink^®^ in situ proximity ligation assay (PLA) system (Sigma-Aldrich) according to the manufacturer’s instructions. Images were acquired using a laser-scanning confocal microscope (LSM510 Meta, Carl Zeiss) equipped with an Axiovert 200M inverted microscope, using a Plan Apo 63/1.4-N oil immersion objective and analyzed with the imaging software Icy. The number of PLA foci per cell was scored manually using the same thresholding parameters across parallel samples.

### Cellular dNTPs quantification by primer extension assay

Differentiated THP1 or U937 cell lines were harvested in ice-cold 65% methanol, lysed (95°C, 3’) and dried using a vacuum concentrator. Dried samples were analyzed for dNTP content were quantified by single nucleotide incorporation assay as described previously^9^.

### Statistical analyses

Graphical representation and statistical analyses were performed using Prism7 software (GraphPad Software, San Diego, CA, USA). Unless otherwise stated, statistical significance was determined by one-way ANOVA test with a Dunnett’s multiple comparison post-test.

### Data availability

All data are available from the corresponding author upon request.

## Acknowledgements

The authors thank H. de Thé and V. Lallemand-Breitenbach for discussion, A. Amara and X. Carnec for critical reading of the manuscript. The authors thank M. Benkirane (IGH, Montpellier, France), N. Manel (I. Curie, Paris, France) and A. Puissant (INSERM U944) for reagents. We are grateful to the Core facility of IRSL and Yasmine Khalil for technical support. This work was supported by Sidaction (grant 2018-1-AEQ-12075 to AZ), Sidaction/FRM (grant VIH2016126003 to AZ), EU FP7 [HEALTH-2012-INNOVATION-1 ‘HIVINNOV’] (Grant no. 305137 to AS and AZ). CM was supported by fellowships from the French “Ministère de la Recherche et de l’Innovation” and Sidaction. Some experiments were performed in the laboratory of B. Kim, supported by NIH grant AI136581 and AI150451 (to BK).

## Author contributions

CM, AC, JTT, AZ conceived and performed most of the experiments; NP performed mutagenesis and cloning; NC, JB and OS planned and performed experiments on primary and iPS-derived macrophages; SAC and BK provided dNTP measurements; MP and FDG contributed to the experiment of Fig. 3C; GB, LE, and FMG provided critical reagents; CM, AC, JTT, AS and AZ interpreted the data; CM, AC, JTT, AZ wrote the original draft; FMG, PL, AZ review and edited the manuscript; AZ supervised the study.

---

**Figure S1.**
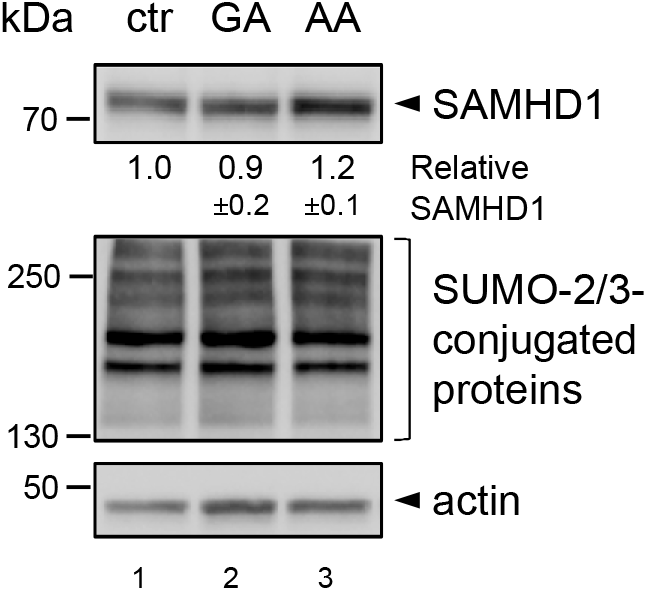
Levels of SAMHD1 are unaffected by inhibition of SUMOylation (related to Fig. 1D and Fig. 6). Proteins (20 μg total proteins/line) contained in the crude lysate of differentiated THP1 treated with GA or AA (50μM, 2h) or DMSO (ctr) were separated by migration on a 4-15% SDS-PAGE gel and, next, visualized by immunoblotting using antibodies against SAMHD1, SUMO2/3, or actin. The intensity of bands corresponding to SAMHD1 and actin, used as loading control, was determined by densitometry with ImageJ software (n=3). The SAMHD1/actin ratio in control cells was set to 1.

**Figure S2.**
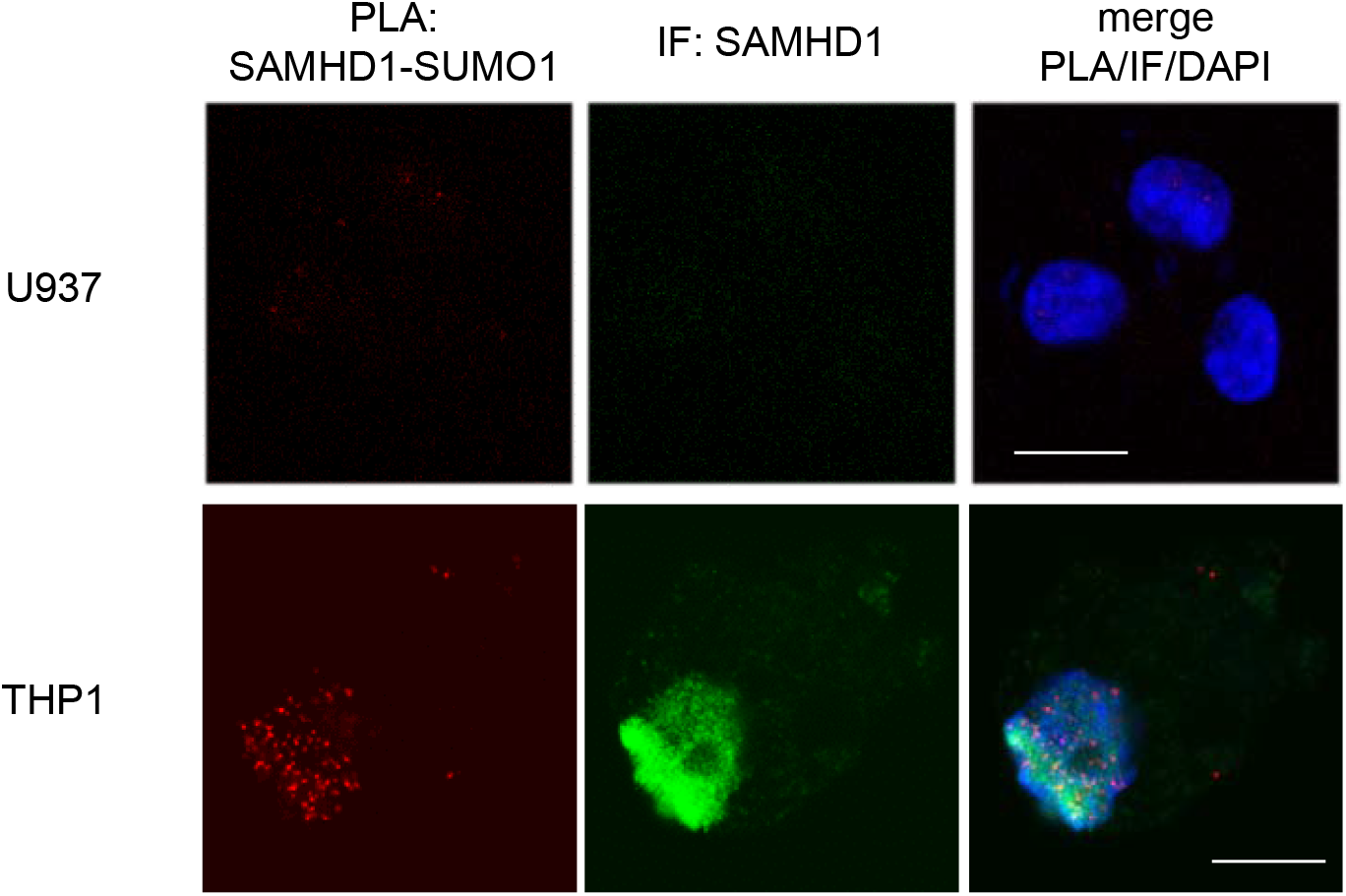
SAMHD1 interacts with SUMO1 in the nucleus of differentiated THP1, but not U937 cells (related to Fig. 1D). Differentiated THP1 or U937 cells were co-stained with anti-SAMHD1 and anti-SUMO1 antibodies and treated as in figure 1D. Data from one representative experiment are shown (n=2). Scale bar = 10 μm.

**Figure S3.**
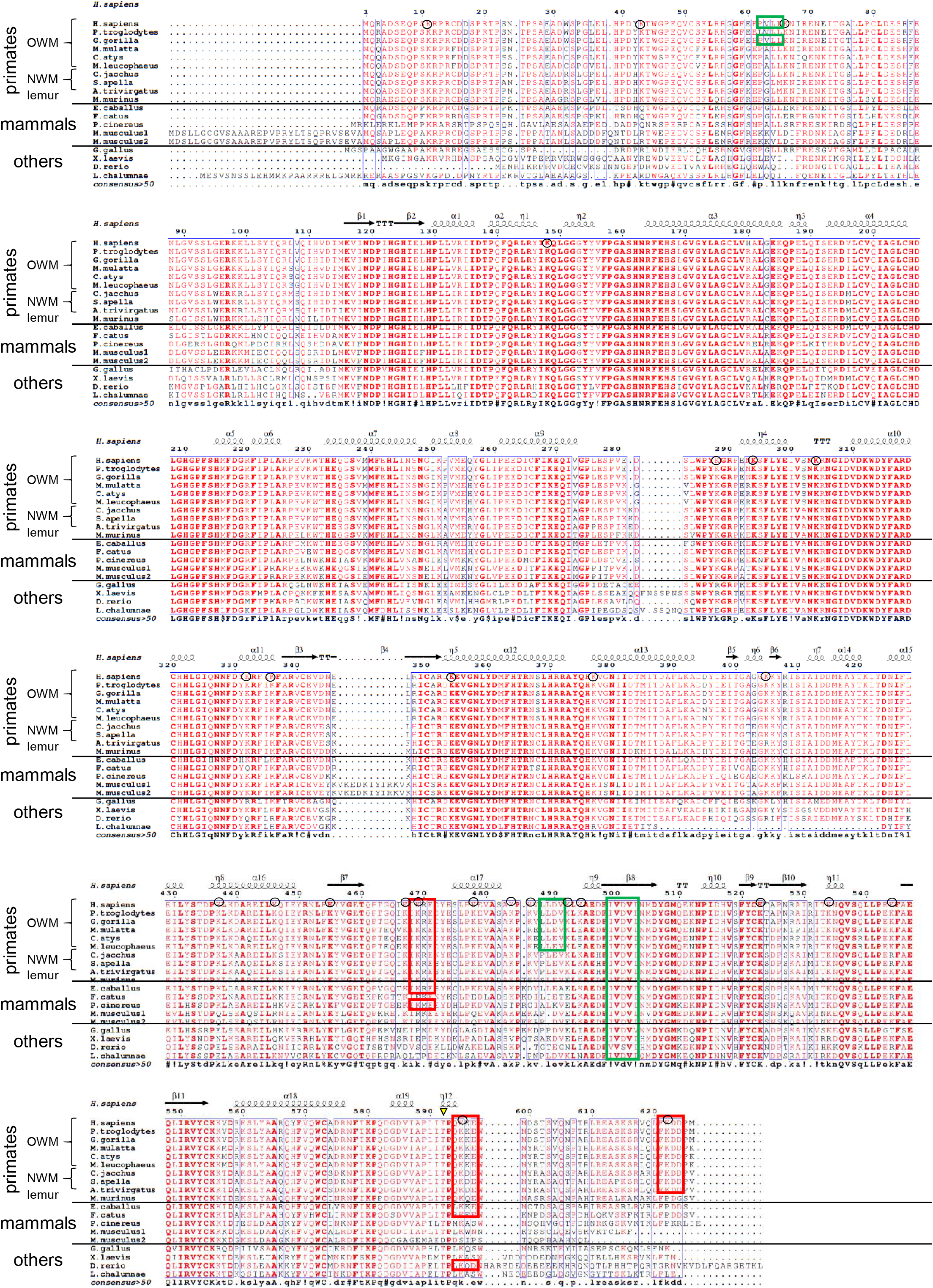
Multi-alignment of SAMHD1 sequences from different species (related to Figs. 2 and 4). Sequences of SAMHD1 isoforms from various vertebrate species were aligned using the MultiAlin software^64^ and visualized using ESPRIT^65^. SUMOylation sites centered on K469, K595 and K622 are boxed in red, while SIM1, SIM2 and SIM3 are boxed in green. The black arrowhead points at the phosphorylatable T592 residue.

**Figure S4.**
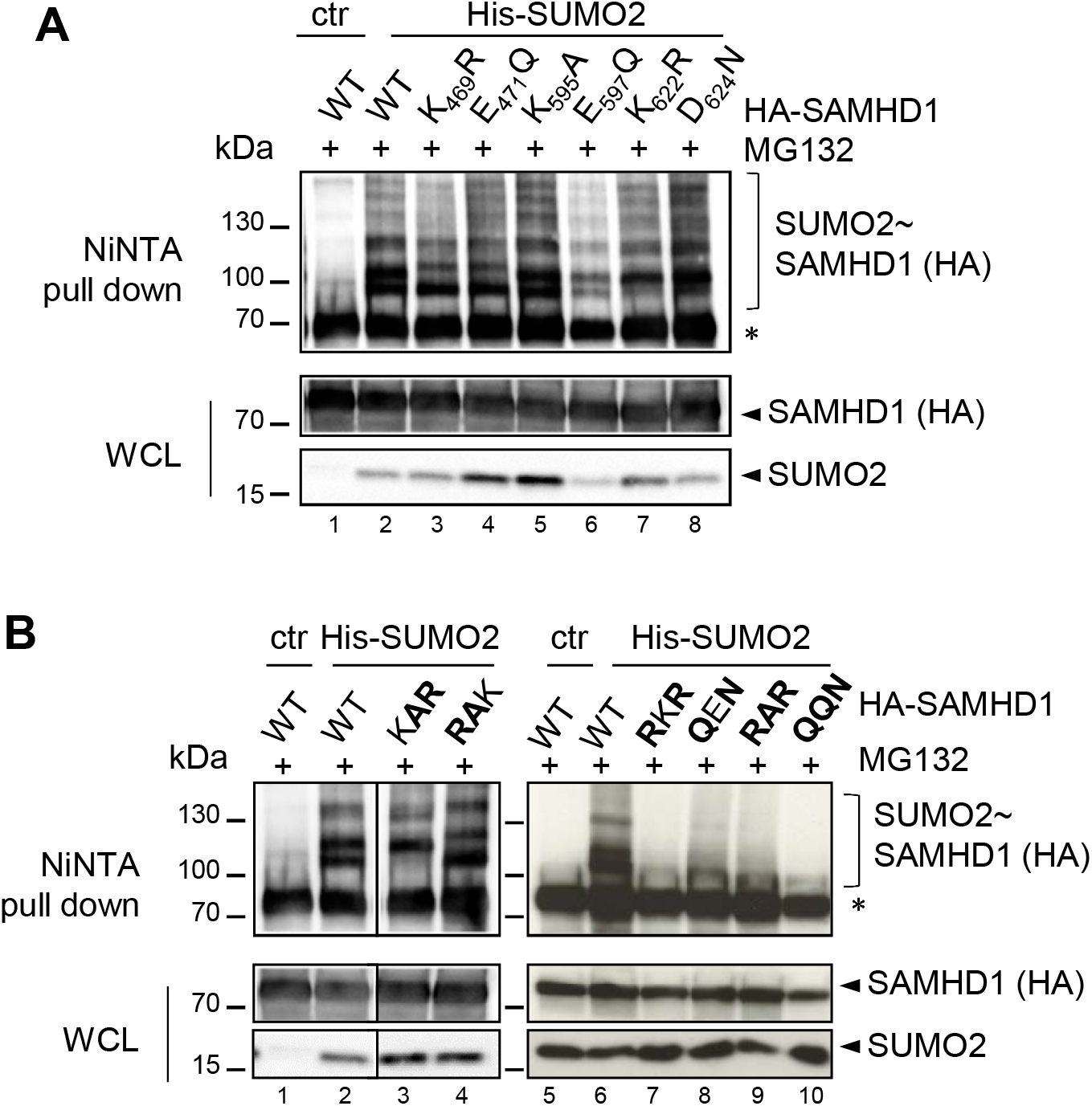
Residues K469 and K622 are modified by SUMO chains that accumulate upon proteasome inhibition (related to Fig. 2). **A.** HEK 293T cells were transfected with plasmids encoding HA-SAMHD1 WT or single or **B.** multiple SUMO-site mutants, Ubc9 and His-SUMO2 or the control empty plasmid (ctr). After 24 hours, samples were treated MG132 (3μM, ON) and then were processed as in Fig. 1A. Bold characters indicate the mutated residues as in Fig. 2C. WCL: whole cell lysate. *, nonspecific binding of unmodified SAMHD1 on Ni-NTA beads. Results of one representative experiment are shown (n≥2).

**Figure S5.**
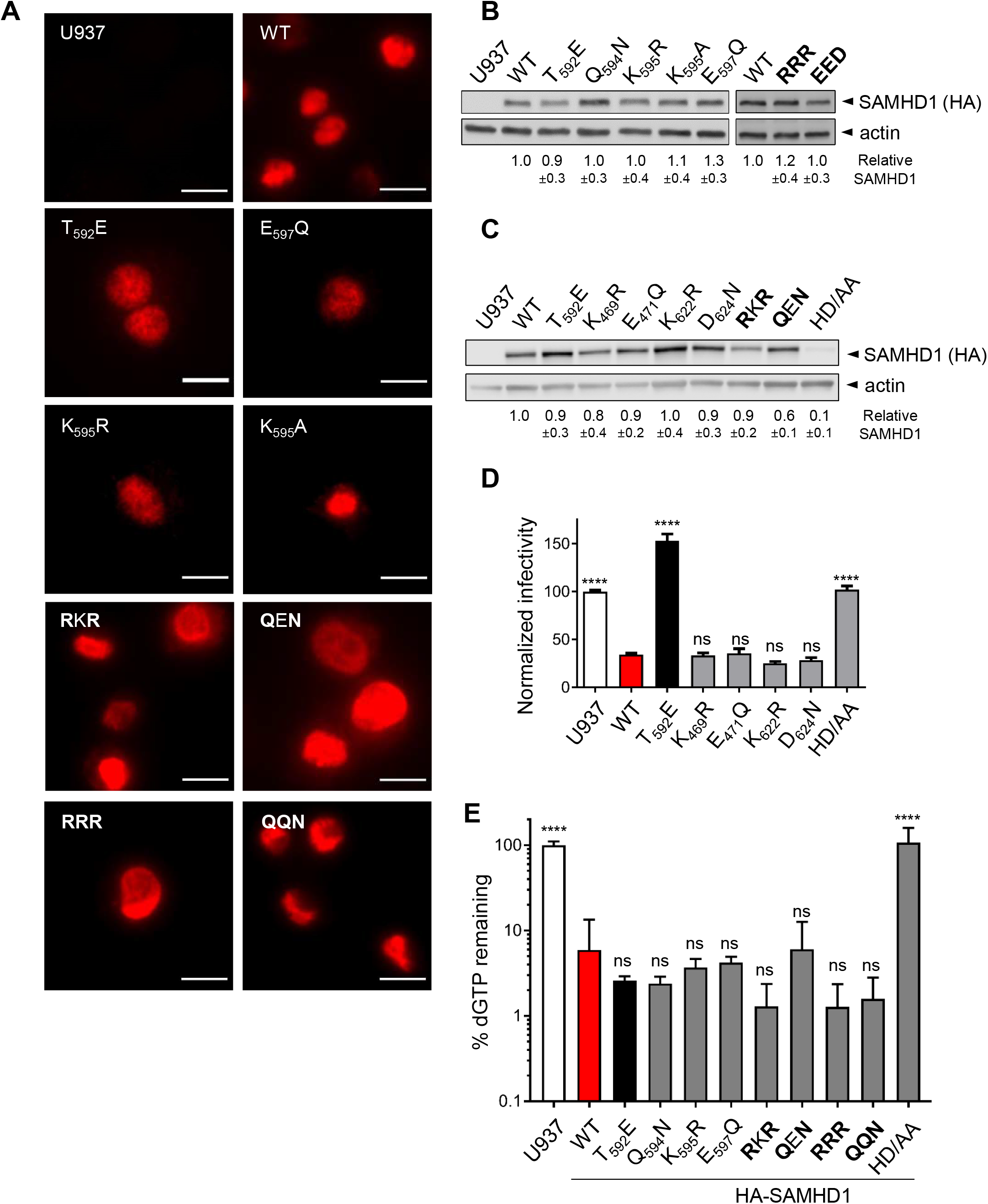
Characterization of SUMOylation-defective SAMHD1 mutants (related to Fig. 3). **A.** The localization of SAMHD1 mutants was assessed in differentiated U937 cells by immunofluorescence using an anti-HA antibody followed by anti-isotype secondary antibody coupled to the Alexa_594_ dye. Images were acquired with an Axiovert 200 M inverted microscope, using a Plan Apo 63/1.4-N oil immersion objective. Scale bar =10 μm. **B.** The expression levels of SAMHD1 single or **C.** multiple SUMO-site variants stably expressed in U937 cells were monitored after PMA treatment (100 μg/mL, 24h) in the total cell extract by immunoblotting (20 μg total proteins/lane). Band intensities were quantified as in Fig. 5A (n=4). The WT SAMHD1/actin ratio was set to 1. Bold characters indicate the mutated residues as in Fig. 2C. **D.** U937 cell lines expressing the indicated HA-SAMHD1 variants (2 independently generated cell lines) were infected with VSVg/HIV-1ΔEnvGFP virus in 4 technical replicates and analyzed as in Fig. 3B. Bars represent the mean ±SD (n=3). Statistical significance was assessed by one-way ANOVA test. ****: p<0.0001. ns: not significant. **E.** Cellular dGTP levels were quantified as previously described^9^. The dGTP levels (%) of SAMHD1-expressing cells were calculated relative to those of parental U937 set to 100. Bars show the mean ± SD (n≥3). Statistical significance was assessed by one-way ANOVA test. ****: p<0.0001. ns: not significant.

**Figure S6.**
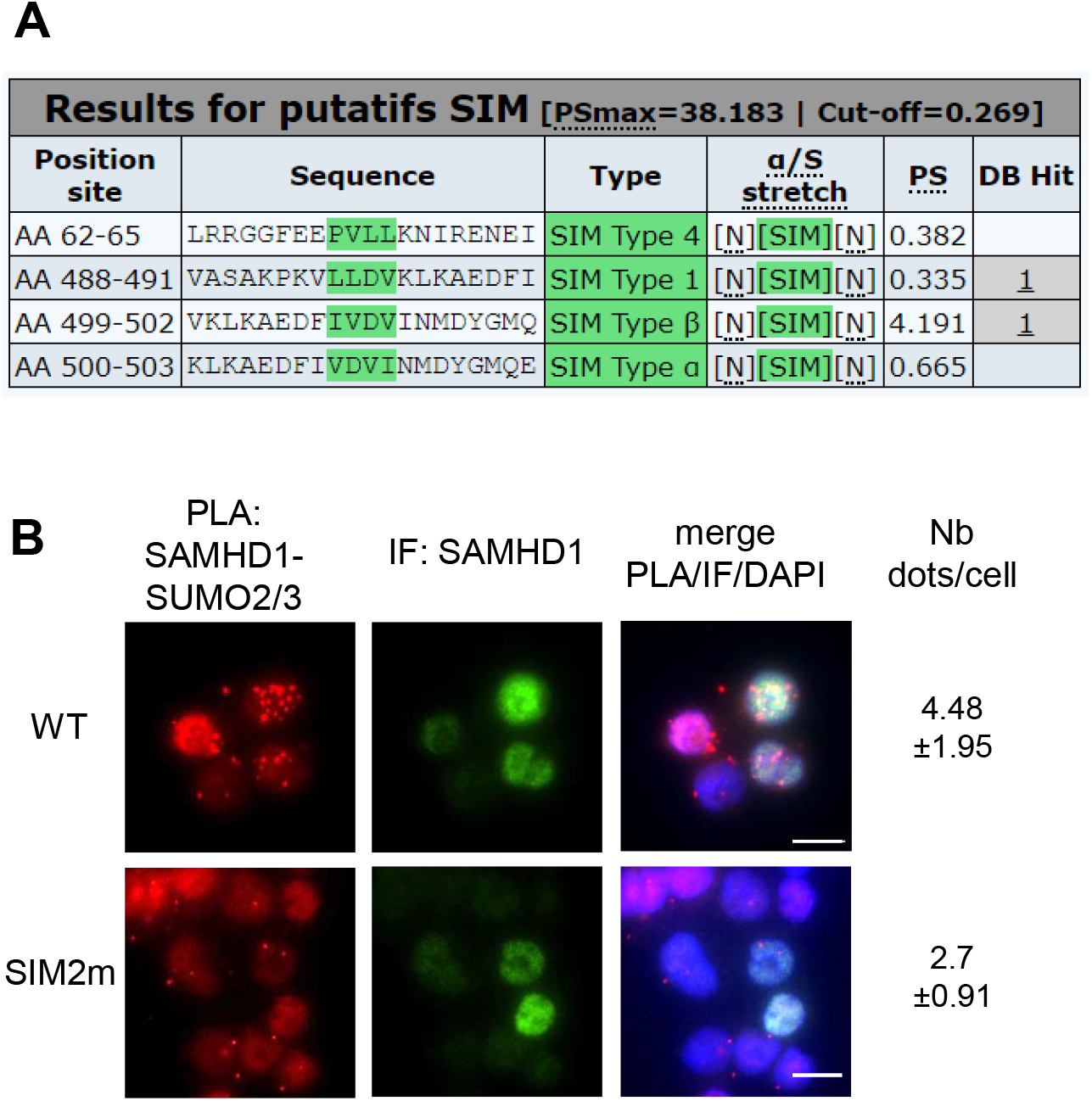
The SIM2 of human SAMHD1 is important for the non-covalent interaction with SUMO proteins (related to Fig. 4). **A.** *In silico* analysis with JASSA^46^ predicts the presence of several SIMs in the sequence of human SAMHD1. **B.** Differentiated U937 cell lines stably expressing WT or SIM2m SAMHD1 variants were processed for PLA and analyzed as in Fig. 1D.

**Figure S7.**
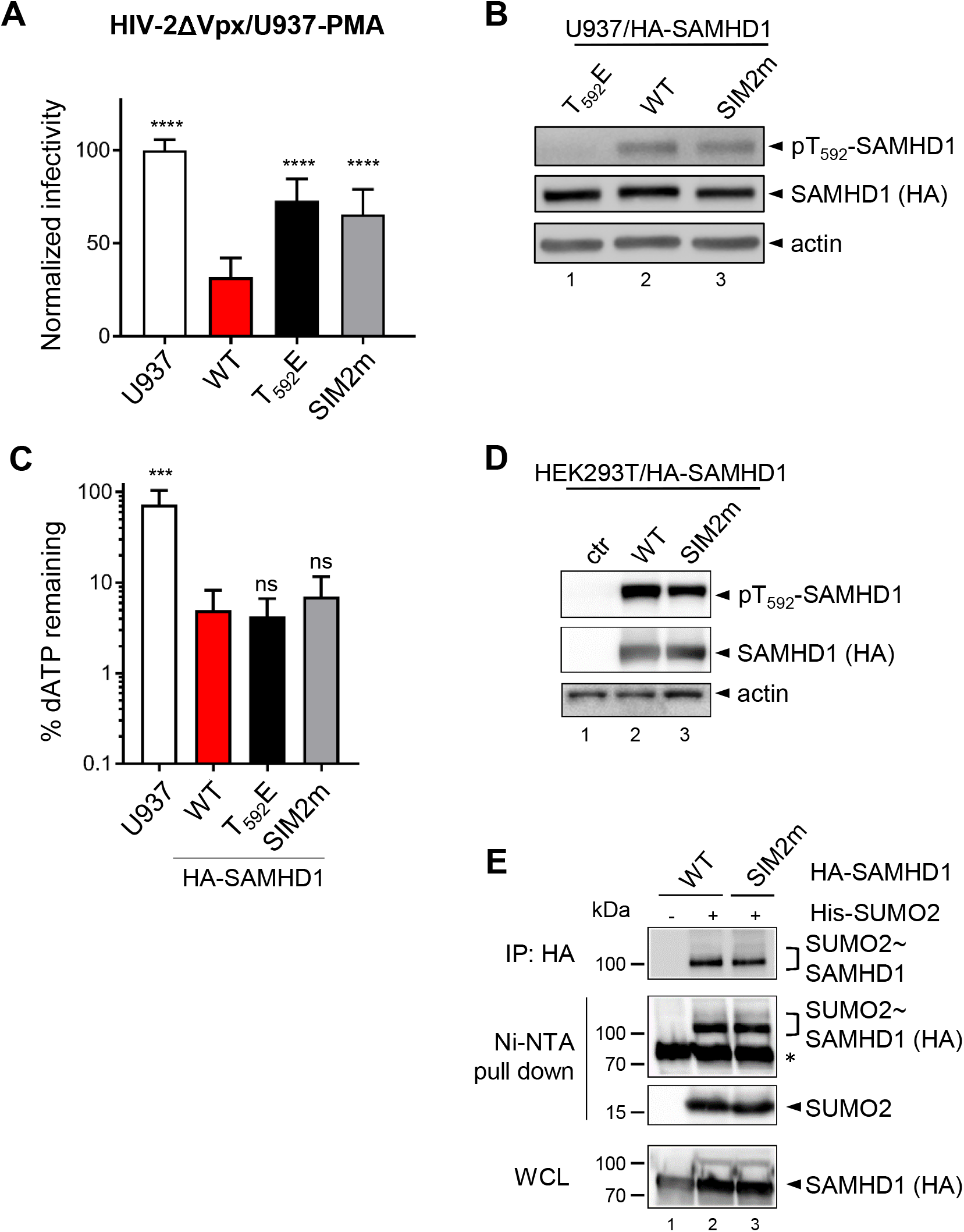
SAMHD1 SIM2m variant does not restrict HIV-2ΔVpx but has WT dNTPase activity (related to Fig. 4). **A.** U937 cells stably expressing HA-SAMHD1 WT or SIM2m mutant (3 independent transductions) were differentiated by PMA treatment (100ng/mL, 24 h) and then challenged with VSVg/HIV-2ΔVpxGFP in 4 technical replicates. Analysis was performed as in Fig. 3B. Bars represent the mean ±SD (n=4). Statistical significance was assessed by one-way ANOVA test. ****: p<0.0001. ns: not significant. **B.** The levels of total and phosphorylated SAMHD1 were monitored in the crude extract of differentiated U937 cell lines using an anti-HA or anti-pT592 specific antibody (10 μg total proteins/line). Actin was used as loading control. The band intensities were quantified by densitometry with ImageJ software (n=3). The SAMHD1/actin and pT592/unmodified SAMHD1 ratios for WT SAMHD1-expressing cells were set to 1. **C.** The levels of dATP were quantified and normalized as in Fig. 3D. The dNTP levels of U937 were set to 100%. Bars show the mean ± SD (n=4). Statistical significance was assessed by one-way ANOVA test. ****: p<0.0001. ns: not significant. **C.** The levels of total and phosphorylated SAMHD1 were monitored in the crude extract of transfected 293T cells and analyzed ad in B. **E.** HEK 293T cells overexpressing WT or SIM2m HA-SAMHD1 mutants, Ubc9 and His-SUMO2 were treated as in Fig. 6E. Results of one representative experiment are shown (n=2). *, nonspecific binding of unmodified SAMHD1 on Ni-NTA beads.

**Figure S8.**
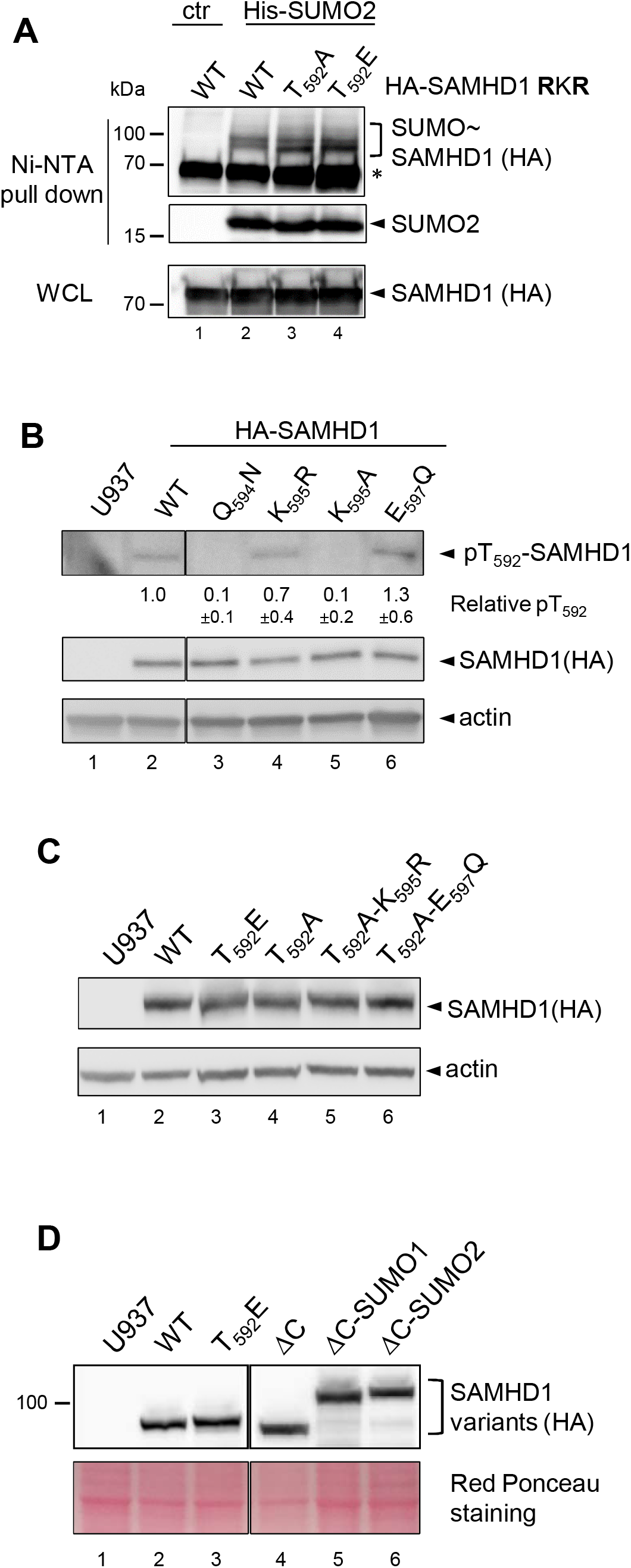
The phosphorylation status of T592 does not influence K595 SUMOylation, while SUMOylation-deficient K595A SAMHD1 mutant is hypophosphorylated (related to Fig. 5). **A.** HEK 293T cells overexpressing the indicated HA-SAMHD1 variants, Ubc9 and His-SUMO2 were processed as in Fig. 1A. WCL: whole cell lysate. One representative experiment is shown (n=4). **B.** The expression levels of SAMHD1 variants stably expressed in differentiated U937 cells were monitored in the total cell extract by immunoblotting (20 μg total proteins/lane). Band intensities were quantified as in Fig. 5A. The WT SAMHD1/actin ratio was set to 1 (n=3). **C. and D.** The expression levels of SAMHD1 variants were analyzed in the crude lysate of differentiation of U937 cell lines (15 μg total proteins/lane).

**Figure S9.**
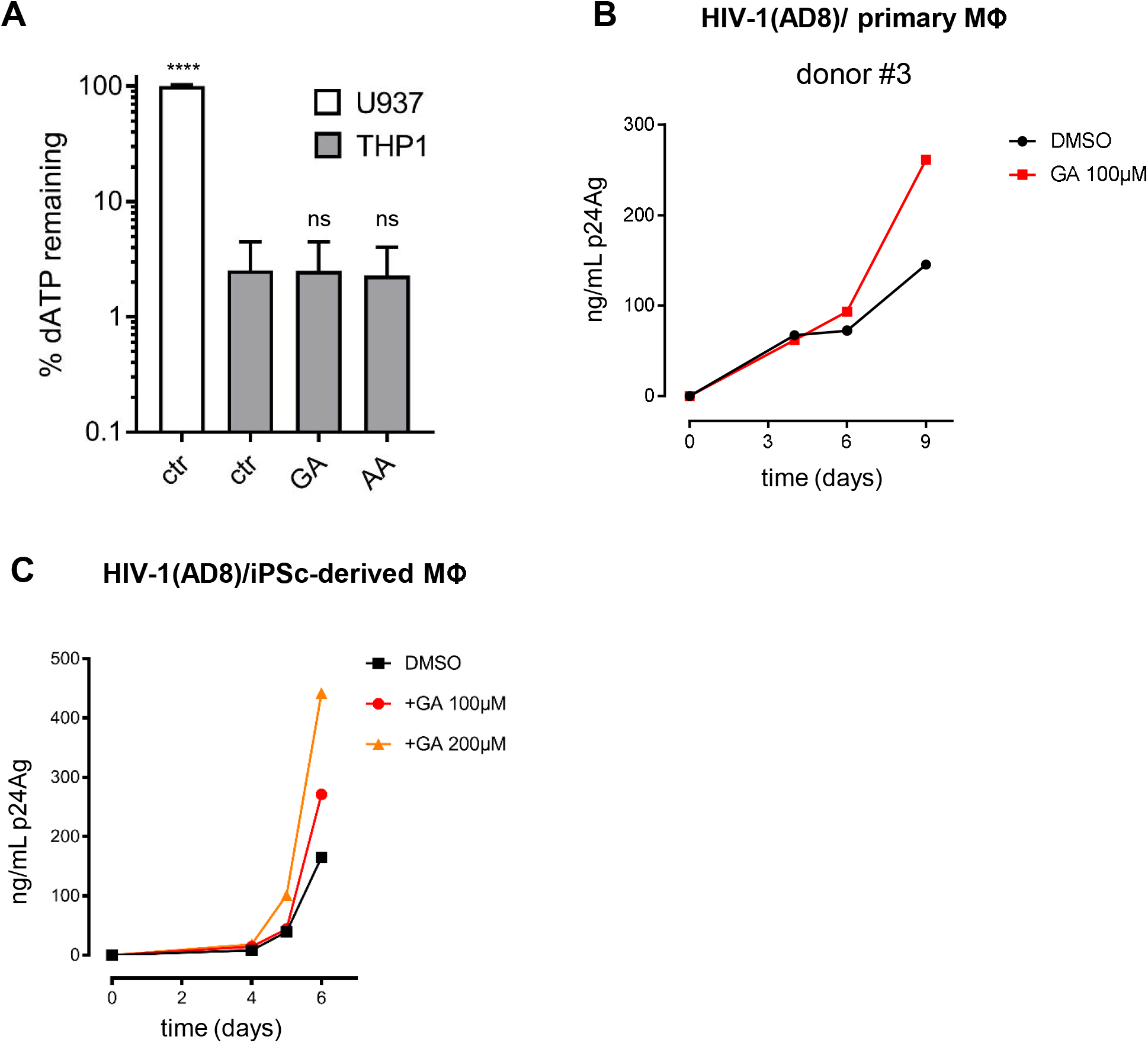
(related to Fig. 6). Inhibition of SUMOylation enhances HIV-1 spreading in human monocyte-derived macrophages without modifying dATP levels. **A.** Cellular dATP levels were quantified as in Figure 3D. The dNTP levels of U937 were set to 100 %. Bars show the mean ± SD (n=4). Statistical significance was assessed by one-way ANOVA test. ****: p<0.0001. ns: not significant. **B.** Viral replication kinetics in primary MDMs from a healthy donor or **C.** macrophages derived from induced-pluripotent stem cells (iPSc) pretreated with gingkolic acid (GA, 2h) and then challenged with the AD8 HIV-1 strain (10 ng/mL p24Ag). Viral replication was analyzed as in Fig. 6C.

