## Supplemental Table 1 for "SUMOylation of SAMHD1 at Lysine 595 is required for HIV-1 restriction in non-cycling cells"

**Table S1. Compilation of SAMHD1 residues identified as potential SUMOylation target sites in large-scale proteomic studies.**

|  | 1 | 2 | 3 | 4 | 5 | 6 | 7 | 8 | 9 | 10 | 11 | 12 | 13 | 14 | 15 | 16 | 17 | 18 | 19 | 20 | 21 | 22 | 23 | 24 | 25 | 26 | 27 | 28 |
| --- | --- | --- | --- | --- | --- | --- | --- | --- | --- | --- | --- | --- | --- | --- | --- | --- | --- | --- | --- | --- | --- | --- | --- | --- | --- | --- | --- | --- |
| **Position (Human SAMHD1)** | **11** | **43** | **66** | **148** | **288** | **294** | **304** | **332** | **336** | **354** | **377** | **405** | **437** | **446** | **455** | **467** | **469** | **478** | **484** | **486** | **492** | **494** | **494** | **523** | **534** | **544** | **595** | **622** |
| **Study count** | **1** | **2** | **2** | **1** | **1** | **2** | **1** | **1** | **1** | **1** | **1** | **1** | **1** | **2** | **2** | **3** | **7** | **1** | **1** | **1** | **2** | **2** | **1** | **2** | **2** | **3** | **5** | **6** |
| Hendriks & Nielsen et al. (2018) |  | **+** |  |  |  | **+** |  |  |  |  |  |  |  |  |  | **+** | **+** |  |  |  |  |  |  | **+** | **+** | **+** | **+** |  |
| Matic & Vertegaal et al. (2010) |  |  |  |  |  |  |  |  |  |  |  |  |  |  |  |  |  |  |  |  |  |  |  |  |  |  |  |  |
| Schimmel & Vertegaal et al. (2014) |  |  |  |  |  |  |  |  |  |  |  |  |  |  |  |  |  |  |  |  |  |  |  |  |  |  |  |  |
| Tammsalu & Hay et al. (2014) / HS |  |  |  |  |  |  |  |  |  |  |  |  |  |  |  |  | + |  |  |  |  |  |  |  |  |  | + | + |
| Impens & Ribet et al. (2014) |  |  |  |  |  |  |  |  |  |  |  |  |  |  |  |  |  |  |  |  |  |  |  |  |  |  |  |  |
| Hendriks & Vertegaal et al. (2014) |  |  |  |  |  |  |  |  |  |  |  |  |  |  |  |  | + |  |  |  |  |  |  |  |  |  |  | + |
| Lamoliatte & Thibault et al. (2014) / MG |  |  |  |  |  |  |  |  |  |  |  |  |  |  |  |  | + |  |  |  |  |  |  |  |  |  |  | + |
| Xiao & Vertegaal et al. (2015) |  |  |  |  |  |  |  |  |  |  |  |  |  |  |  |  |  |  |  |  |  |  |  |  |  |  |  |  |
| Hendriks & Vertegaal et al. (2015-CR) |  |  |  |  |  |  |  |  |  |  |  |  |  |  |  |  |  |  |  |  |  |  |  |  |  |  |  |  |
| Hendriks & Vertegaal et al. (2015-NC) |  |  |  |  |  |  |  |  |  |  |  |  |  |  |  |  |  |  |  |  |  |  |  |  |  |  |  |  |
| Lamoliatte & Thibault et al. (2017) / MG |  |  | + | + |  |  |  |  |  |  |  |  |  | + | + | + | + |  | + |  | + | + |  |  | + | + | + | + |
| Hendriks & Nielsen et al. (2017) | + | + | + |  | + | + | + |  | + | + | + | + | + | + | + | + | + |  |  | + | + | + | + | + |  | + | + | + |
| Lumpkin & Komives et al. (2017) |  |  |  |  |  |  |  | + |  |  |  |  |  |  |  |  | + | + |  |  |  |  |  |  |  |  | + | + |
